# Structural basis of interaction between Pol1 and DnaN from *Mycobacterium tuberculosis*

**DOI:** 10.64898/2026.09.19.751633

**Authors:** Dalchand Sharma, Deepak T. Nair

## Abstract

DNA polymerases generally interact with the processivity clamp (DnaN/β-clamp) during replication via clamp-binding motifs (CBMs). Since a CBM was not detected in DNA Polymerase I (PolI) from *E. coli*, it is generally believed that PolI performs its functions in genome replication and repair, independently of the β-clamp. We observe that, while PolI and β-clamp proteins from *E. coli* do not interact, their homologs from *Mycobacterium tuberculosis* bind strongly, and the binding site localises to the Klenow fragment of PolI. The cryo-EM structure (4.9 Å) of the Mtb*-*PolI-Klenow: Mtb-β-clamp complex, coupled with continuous heterogeneity analysis of the cryo-EM data and binding experiments, showed that the primary CBM in PolI may be defined by the stretch ^457^QLSLLDD^463^. The crystal structure (2.9 Å) of the β-clamp with the CBM peptide revealed specific interactions that stabilise the peptide within the β-clamp’s canonical binding site. Phylogenetic analyses across different bacteria showed that the identified CBM is present only in PolI from members of the order Mycobacteriales. Overall, the interaction between PolI with the β-clamp, mediated by a CBM unique to the Mycobacteriales order, highlights the mechanistic distinctiveness of genome replication in Mtb and phylogenetically related organisms, and has important implications for drug discovery.

## Introduction

DNA polymerases (dPols) are centrally involved in DNA synthesis during genome replication, repair and recombination^1,2^. In prokaryotes, the duplication of the bacterial chromosome is generally carried out by DNA polymerase III (PolIII), which belongs to the C-family of dPols^3,4^. The other essential dPol is DNA polymerase I (PolI), a product of the *polA* gene and a member of the A-family of polymerases^3^. PolI usually possesses three activities, 5’ to 3’ exonuclease (EXO1), 5’ to 3’ polymerase (POL), and 3’ to 5’ exonuclease (EXO2) ^5^. During bacterial replication, PolI is involved in the maturation of Okazaki fragments to complete the synthesis of the lagging strand. The POL and EXO2 activities are responsible for high-fidelity polymerisation and proofreading, respectively, and together constitute the Klenow fragment. The 5’-3’ flap endonuclease by the EXO1 domain removes RNA primers and aids nick translation. PolI from *M. smegmatis* may also possess reverse transcriptase activity and perform RNA/DNA flap-nick translation activity^5^. Mtb-PolI (Rv1629) naturally lacks 3’-5’ proofreading activity due to the absence of metal-binding amino acid residues in the active site^6^. The disruption of the polymerase region of *polA* gene in the case of *M. smegmatis* led to hypersensitivity towards DNA-damaging agents^7,8^. The structures of PolI from *E. coli* (Klenow) and *M. Smegmatis* (full-length) provide insight into the architecture of the enzyme^5,9^.

In addition to PolIII and PolI, prokaryotes also possess B-, X-and Y-family dPols that have specialised functions during replication and repair^10–14^. The dPols in *E. coli* are generally believed to interact with the processivity clamp during replication and repair^15–18^. This protein, a product of the *dnan* gene, is also known as the β-subunit (of PolIII holoenzyme) or β-clamp. It is a toroidal protein that encircles DNA, increases the processivity of dPols and aids recruitment of proteins involved in repair and replication^19,20^. The interaction of the dPols with the clamp generally occurs through a clamp-binding motif (CBM), which binds to the canonical binding site on the clamp^19^. Although some studies suggest that the β-clamp binds to PolI in *E. coli* and influences its activity, the region of the enzyme that mediates these interactions has not yet been identified, leading to the hypothesis that PolI’s functions may occur independently of the clamp^21–23^.

We observe that, unlike the proteins from *E. coli*, the β-clamp and PolI from *M. tuberculosis* form a strong interaction and that the interaction sites are located within the Klenow fragment of Mtb-PolI. Using a combination of structural, computational and biochemical tools, we have identified the CBM of PolI that interacts with the canonical binding site on the β-clamp. Phylogenetic analysis of PolI sequences from different bacteria revealed that the peptide motif is present only in members classified in the order Mycobacteriales within the Actinomycota class. Overall, our studies have identified and characterised the CBM present in PolI from bacteria in the Mycobacteriales order, and this discovery has implications for genome replication in organisms in this order and for drug discovery.

## Results

### PolI and β-clamp from *Mycobacterium tuberculosis* interact with high affinity

Microscale Thermophoresis (MST) experiments showed that Mtb-PolI-Klenow and Mtb-PolI-FL could bind Mtb-β-clamp with K_d_ values of 160 nM and 1100 nM, respectively (Fig. 1A). The Mtb-PolI-EXO1 domain construct showed no binding with the Mtb-β-clamp protein. In the case of *E. coli* proteins, Ec-PolI-Klenow and Ec-PolI-FL bound Ec-β-clamp with K_d_ values of 15.3 µM and 13 µM, respectively (Supplementary Fig. S1A). These experiments showed that Mtb-β-clamp interacts with Mtb-PolI-Klenow with nearly 10-fold higher affinity than with Mtb-PolI-FL. The Ec-PolI-Klenow and Ec-PolI-FL proteins bind to Ec-β-clamp with nearly 80 times lower affinity than the corresponding interaction between Mtb-β-clamp and Mtb-PolI-Klenow. These studies show a strong interaction between Mtb-PolI and Mtb-β-clamp, and that the interacting site in Mtb-PolI is located within the Klenow fragment.

**Figure 1:**
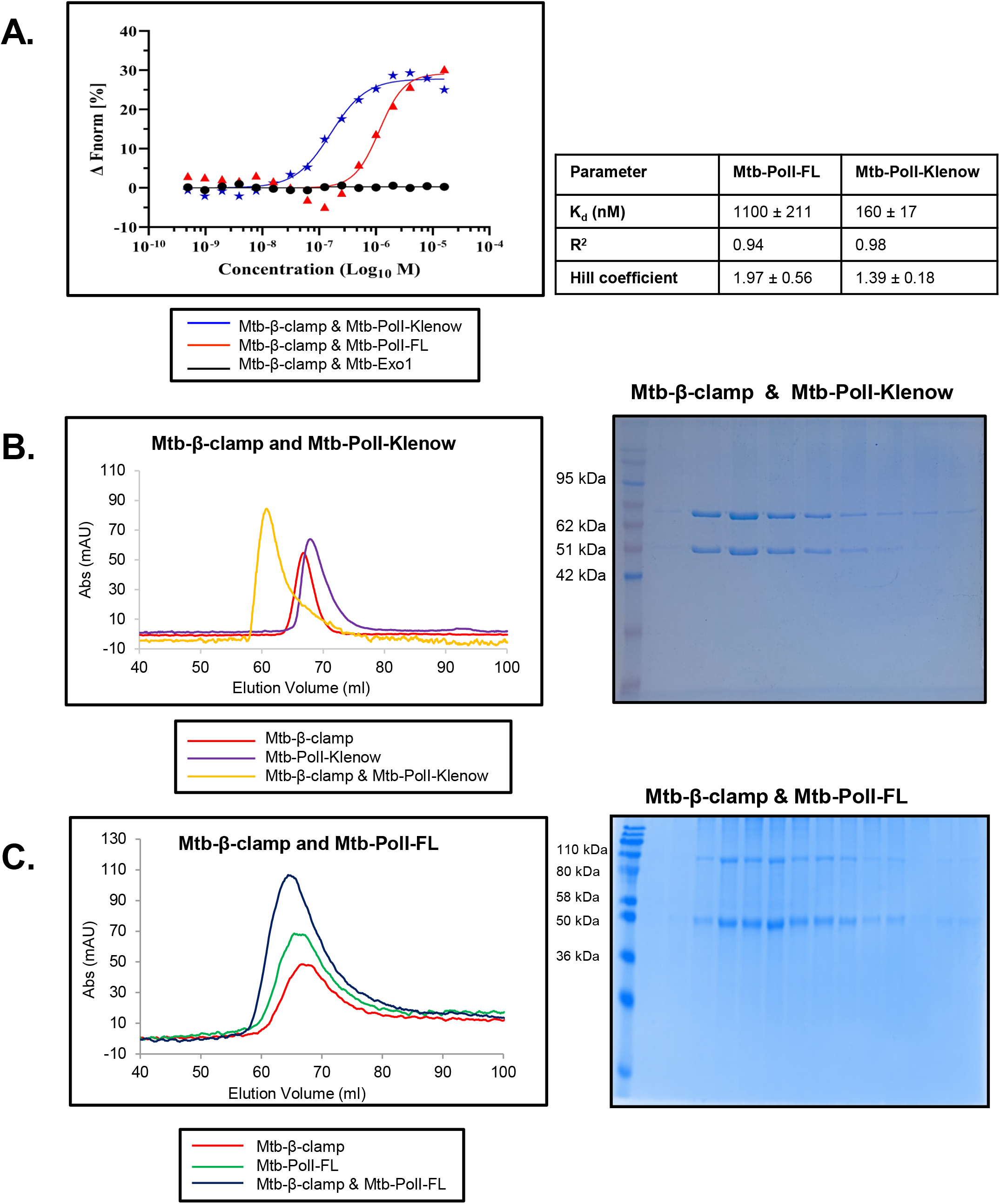
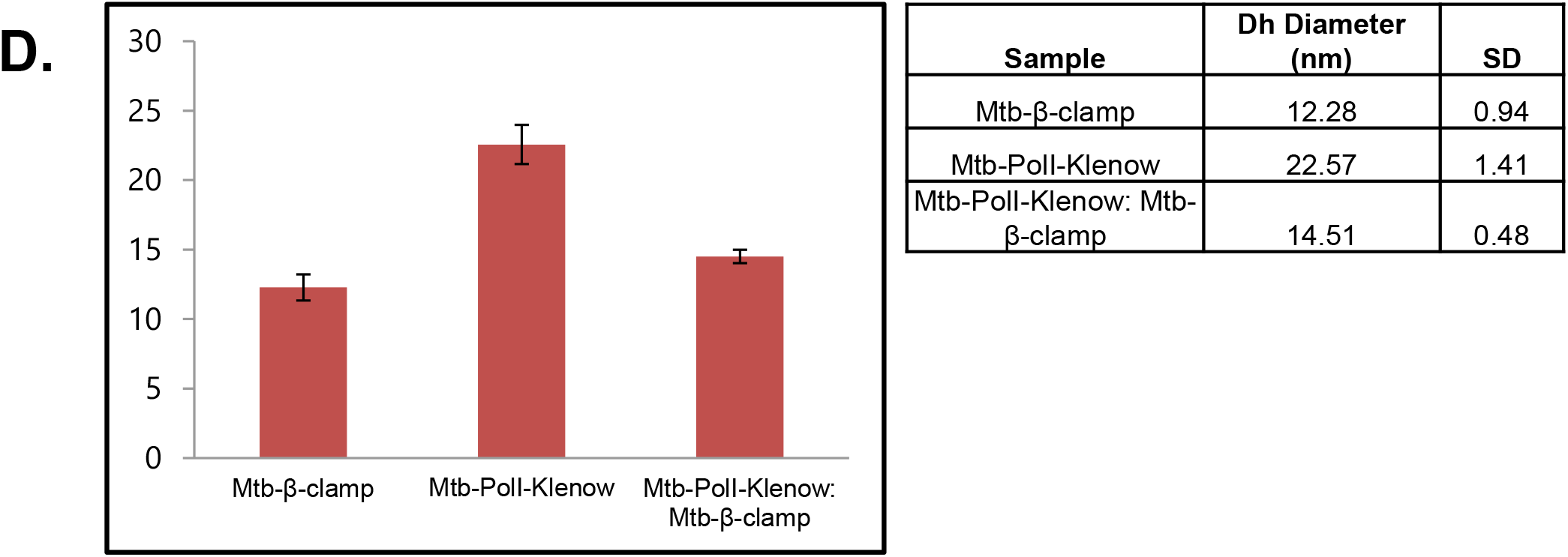
Assessment of the interaction of different constructs of Mtb-PolI with Mtb-β-clamp. (A) Binding curves are displayed for data generated using MST, for the interaction of Mtb-β-clamp with Mtb-PolI-FL (red), Mtb-PolI-EXO1 domain (black) and Mtb-PolI-Klenow (blue). (B) Gel filtration analysis was carried out on Mtb-β-clamp (red), Mtb-PolI-Klenow (purple) and Mtb-β-clamp mixed with Mtb-PolI-Klenow (orange). The corresponding SDS-PAGE analysis of the eluted fractions from the sample containing a mixture of Mtb-β-clamp and Mtb-PolI-Klenow is shown. (C) Gel filtration analysis was carried out on Mtb-β-clamp (red), Mtb-PolI-FL (green) and Mtb-β-clamp mixed with Mtb-PolI-FL (dark blue). The corresponding SDS-PAGE analysis of the eluted fractions from the sample containing a mixture of Mtb-β-clamp and Mtb-PolI-FL is shown. (D) Dynamic Light Scattering (DLS) analysis of the purified Mtb-β-clamp, Mtb-PolI-Klenow and Mtb-PolI-Klenow:Mtb-β-clamp complex.

Gel filtration analysis showed that, when loaded separately, Mtb-β-clamp and Mtb-PolI-Klenow eluted as single peaks at 66.8 ml and 67.9 ml, respectively. For the mixture of Mtb-β-clamp and Mtb-PolI-Klenow, a single peak was observed at a retention volume of 60.8 ml. The elution profile indicated a clear shift in the elution peak in the presence of both proteins. SDS-PAGE analysis showed that the shifted peak contains both proteins (Fig. 1B), and therefore Mtb-β-clamp and Mtb-PolI-Klenow form a stable complex that can be purified by gel filtration. Mtb-PolI-FL eluted at a retention volume of 65.6 ml, and for the mixture of Mtb-β-clamp and Mtb-PolI-FL, a single broad peak at a retention volume of 64.4 ml was obtained (Fig. 1C). A merged elution profile was observed in the presence of both proteins, with no distinct peaks corresponding to the individual proteins or a stable complex of the expected size. The SDS-PAGE analysis shows that the peak contains both proteins, as expected from the merged elution profile.

The homogeneity of the purified Mtb-PolI-Klenow:Mtb-β-clamp complex was confirmed by Dynamic Light Scattering (DLS), which showed a single, sharp peak indicating a homogeneous protein complex. Further analysis showed that the hydrodynamic diameter (nm) of Mtb-β-clamp, Mtb-PolI-Klenow and Mtb-PolI-Klenow:Mtb-β-clamp complex were 12.3 nm, 22.6 nm and 14.5 nm, respectively (Fig. 1D). As the hydrodynamic radius obtained for the Mtb-PolI-Klenow:Mtb-β-clamp was between those obtained for individual proteins, it appears that Mtb-PolI-Klenow adopts a more compact structure on complexation with Mtb-β-clamp.

The corresponding proteins from *E. coli*, Ec-β-clamp, Ec-PolI-FL, and Ec-PolI-Klenow eluted as separate peaks with retention volumes of 67.0 ml, 63.8 ml and 70.4 ml, respectively (Supplementary Fig. S1B & S1C). Unlike the Mtb proteins, the mixture of Ec-β-clamp and Ec-PolI-Klenow yielded separate peaks during gel filtration. Similarly, the mixture of Ec-β-clamp and Ec-PolI-FL yielded separate peaks, and the presence of individual proteins in these peaks was confirmed by SDS-PAGE analysis (Supplementary Fig. S1B & S1C).

### CryoEM structure shows that Mtb-PolI interacts with Mtb-β-clamp through a unique CBM

Since Mtb-β-clamp and Mtb-PolI-Klenow form a stable complex, we collected cryoEM data to determine the structure of this heteromeric complex. Initially, the collected data yielded a map at 7.87 Å resolution (Fig. 2). The previously determined crystal structure of Mtb-β-clamp (3RB9) and a homology model of Mtb-PolI-Klenow could be docked into the map. The structure shows that a region between the vestigial proofreading exonuclease domain and the palm subdomain interacts with the processivity clamp’s canonical binding site. In addition, the structure revealed another interface between residues in the thumb of Mtb-PolI and Mtb-β-clamp. The interacting region is located largely in domain I of the other monomer (B) in the Mtb-β-clamp. Continuous heterogeneity analysis conducted using a 3DFlex workflow (Supplementary Fig. S2) and showed that the thumb could move away from the interface located in domain I of the Mtb-β-clamp (Fig. 3 & Supplementary Movie S1). In contrast, the region between the vestigial proofreading exonuclease domain and the palm subdomain from Mtb-PolI-Klenow remained close to the canonical binding site on the Mtb-β-clamp in the movie generated from the continuous heterogeneity analysis.

**Figure 2:**
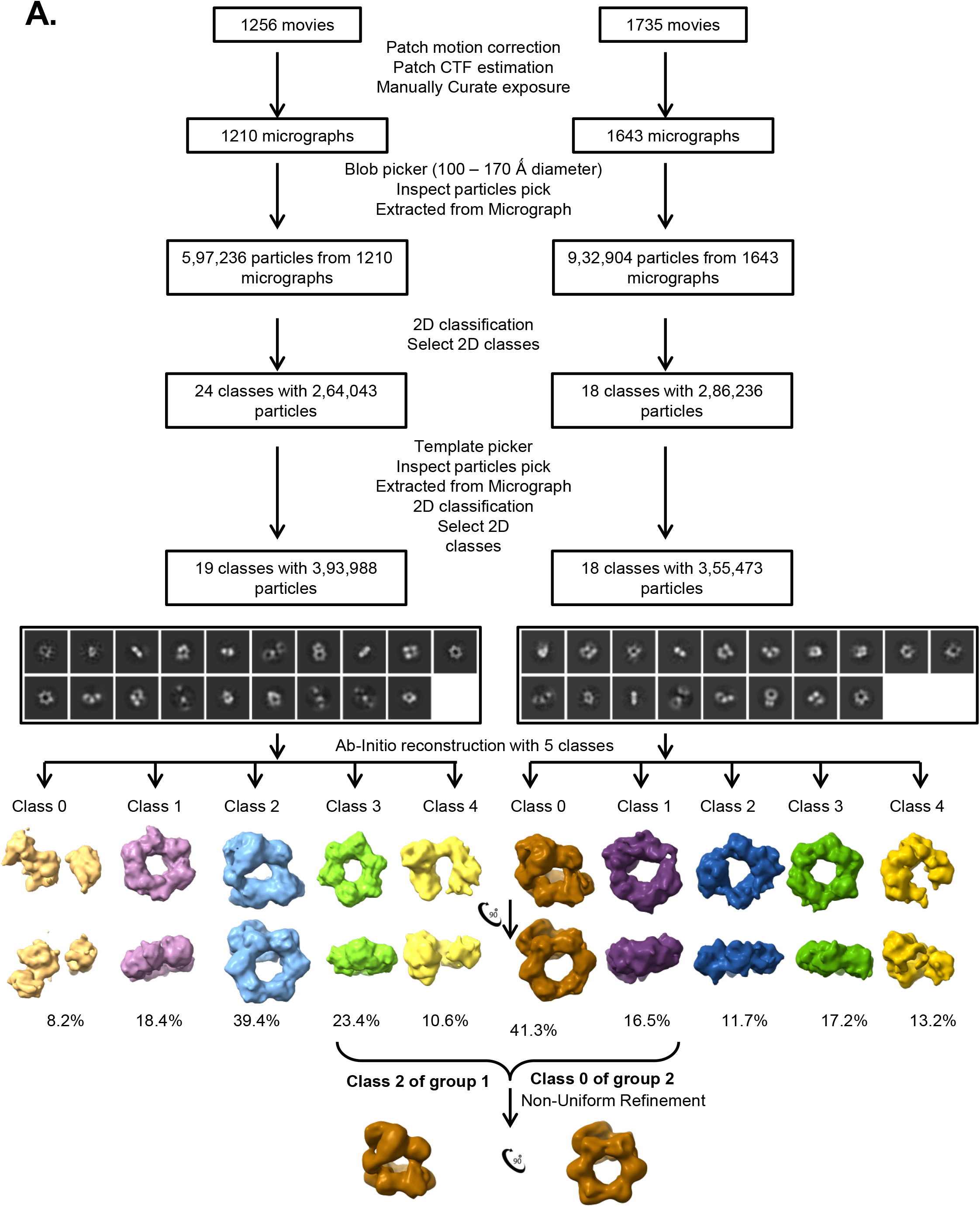
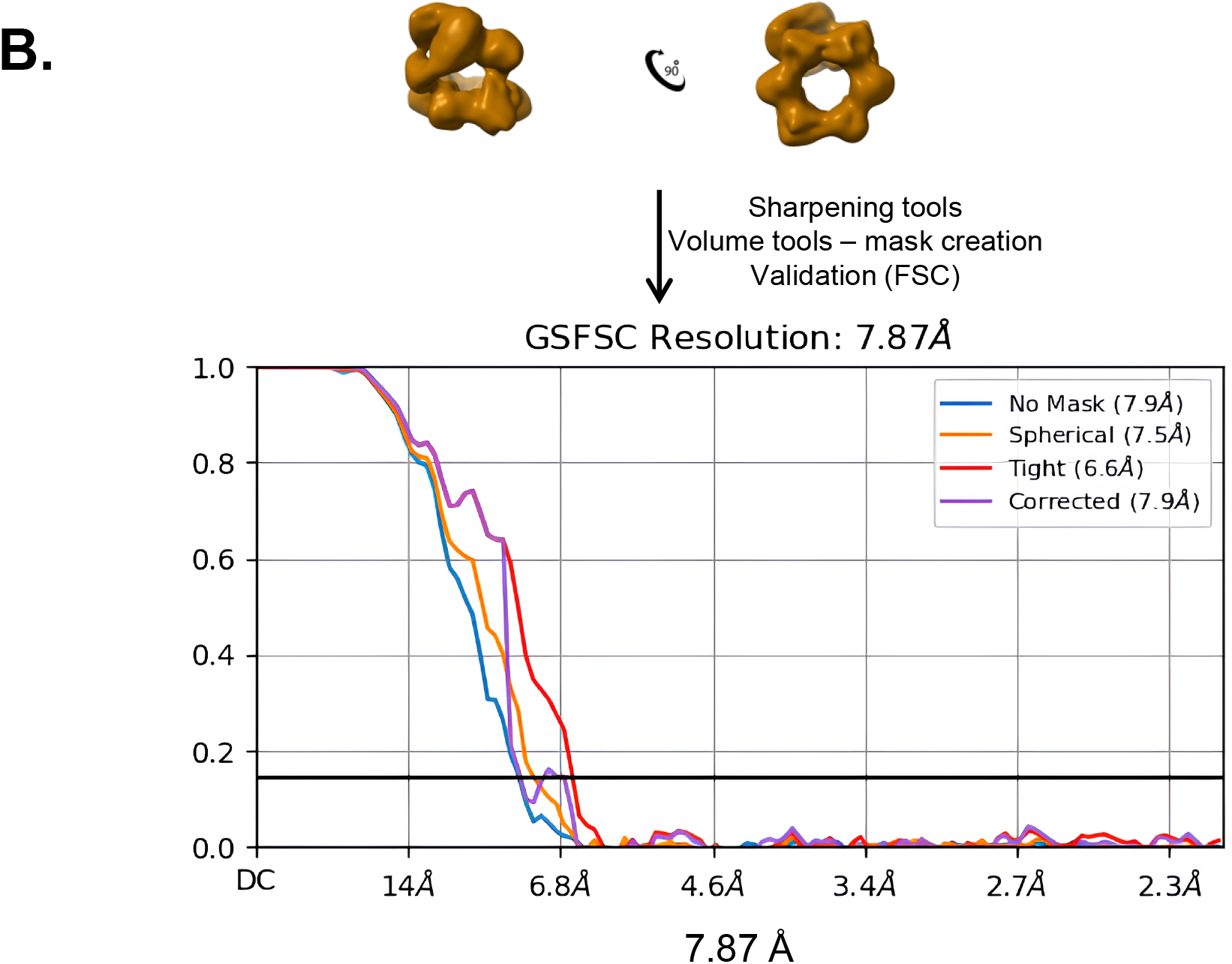
Single particle reconstruction workflow of the Mtb-PolI-Klenow:Mtb-β-clamp complex: (A) Schematic diagram of cryo-EM data processing from two independent datasets using cryoSPARC. Motion correction, CTF estimation, particle picking, and 2D classification were performed, followed by ab-initio reconstruction and heterogeneous refinement. Selected particles from both datasets were combined for the final map reconstruction. Representative 2D class averages, *ab initio* reconstruction classes, and corresponding particle distributions are shown. (B) Gold standard FSC (GSFSC) analysis of final cryo-EM reconstruction of the Mtb-PolI-Klenow: Mtb-β-clamp complex. The final map reached an overall resolution of 7.87 Å following sharpening and local refinement. FSC curves corresponding to masked, unmasked, and masked without noise and phase randomised corrected maps are shown.

**Figure 3:**
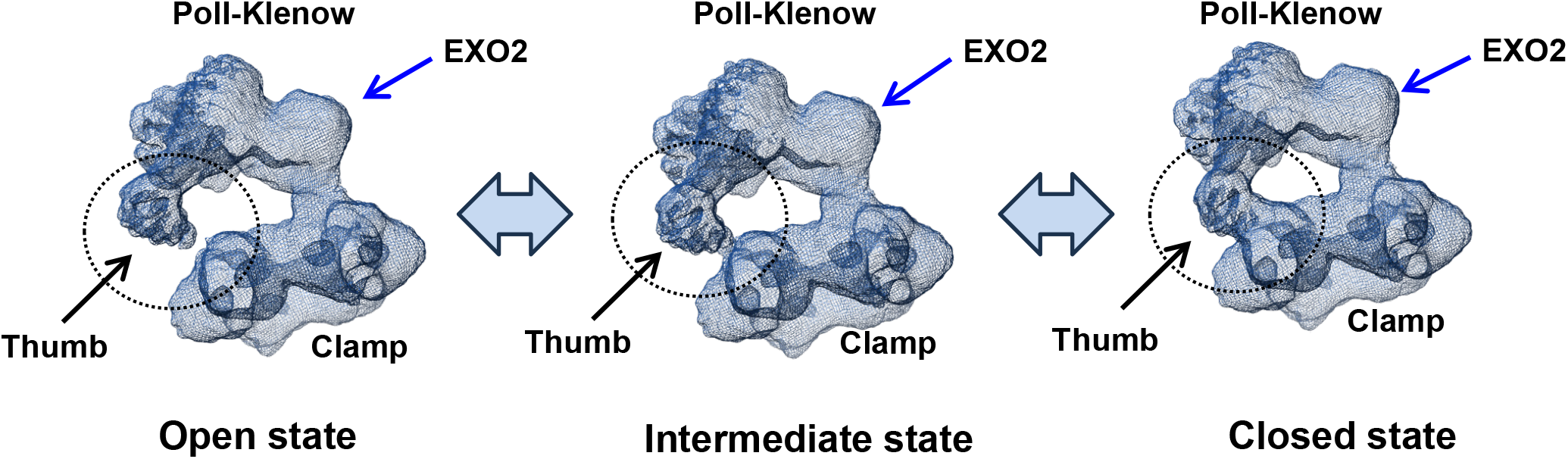
Structural heterogeneity analysis of Mtb-PolI-Klenow:Mtb-β-clamp complex using 3DFlex: Continuous heterogeneity analysis was performed using 3DFlex on low-resolution cryo-EM reconstruction of Mtb-PolI-Klenow:Mtb-β-clamp complex. Three representative snapshots extracted from the series generated by 3DFlex are displayed. Black and blue arrows indicate the thumb subdomain and the EXO2 domain, respectively, and the dotted circle marks the site of dynamic interaction between the thumb subdomain and the clamp.

Using a pipeline combining cryoSPARC and cryoPROS (Supplementary Fig. S3 & Fig. 4), a higher-resolution map with a maximum resolution of 4.9 Å was generated (Supplementary Movie S2). The structure of the complex from the 7.87 Å map was docked into the 4.9 Å map, and manual rebuilding, coupled with real-space refinement, was performed to improve the fit of the complex to the map. Regarding the final structure, 90% of residues are within the density, and the overall Q-value is 0.148. The data collection and validation statistics for the final structure are shown in Table 1. The structure showed that one molecule of Mtb-PolI-Klenow was bound to one dimer of Mtb-β-clamp (Fig. 5 and Supplementary Movie S3). The structure of the Mtb-β-clamp dimer is similar to those deposited in the PDB with accession codes 3P16, 3RB9, 4TR7, 5AGU, 6FVN and 6FVO, and superimposes onto these structures with RMSD values of 1.7 Å, 1.6 Å, 1.8 Å, 1.7 Å, 1.8 Å and 1.7 Å, respectively. The structure of Mtb-PolI-Klenow exhibits a similar backbone structure to that determined for the ortholog from *M. smegmatis,* and the structures superimpose with an RMSD of 2.1 Å.

**Figure 4:**
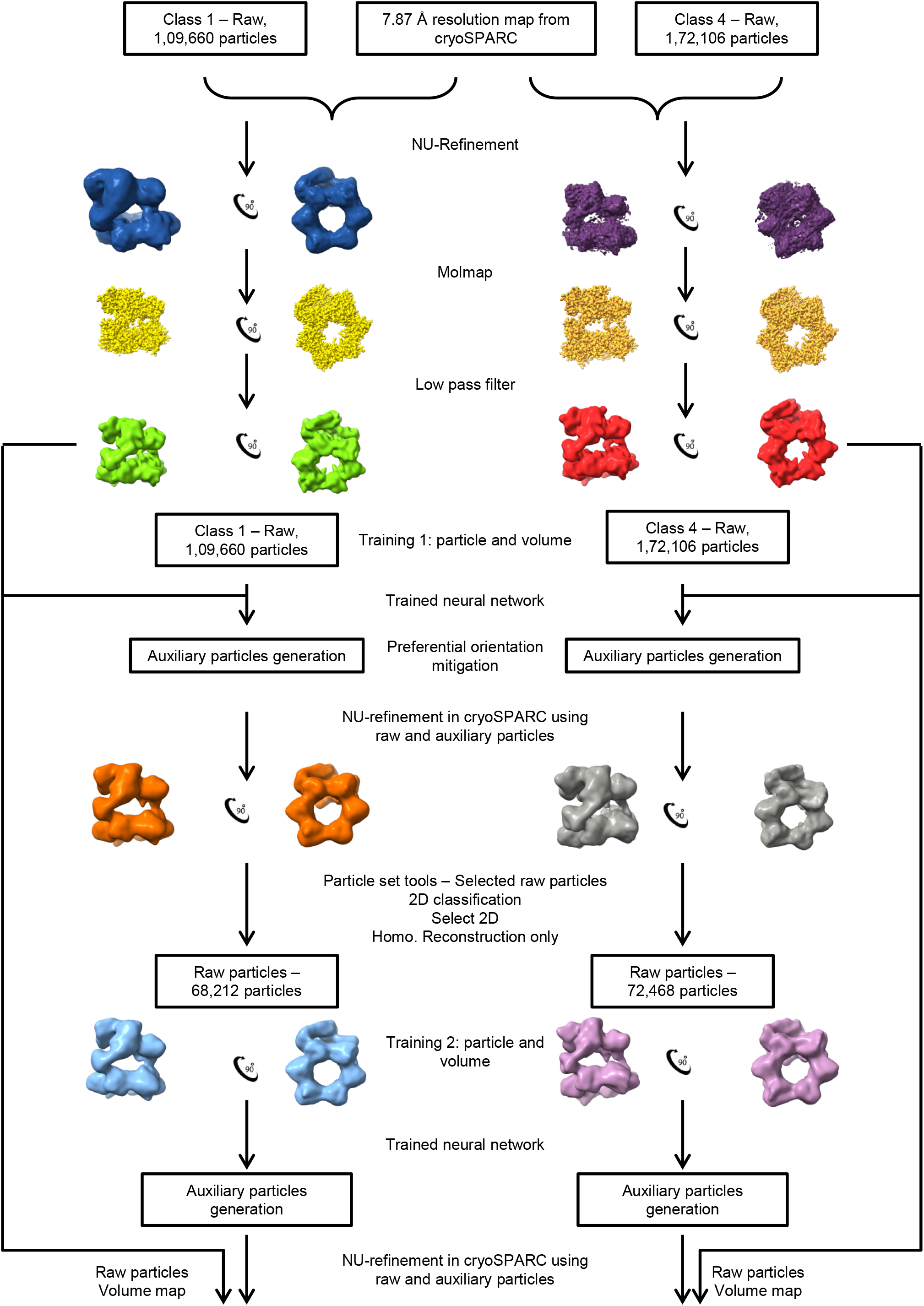
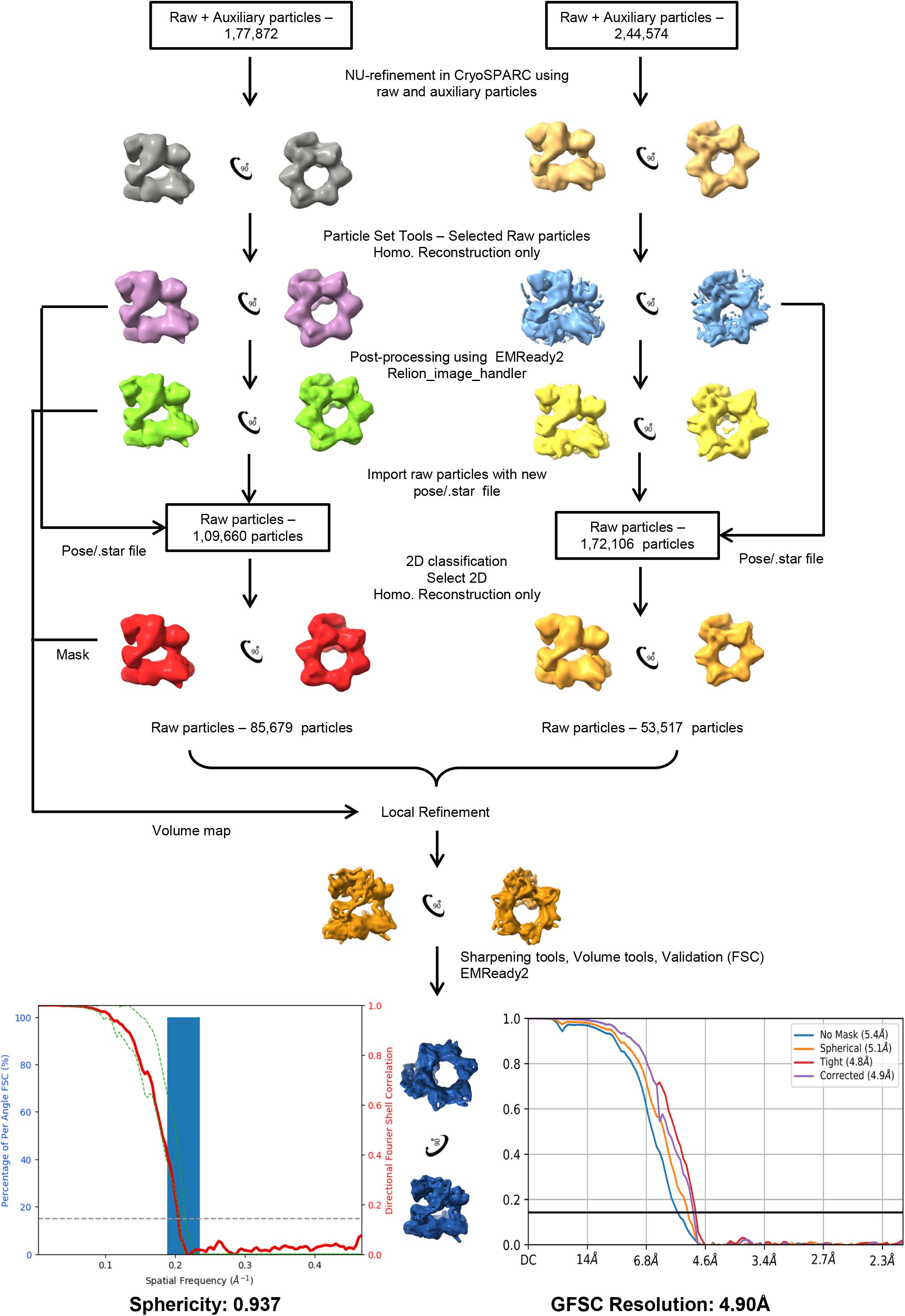
Resolution improvement using cryoPROS-assisted data processing. Schematic representation of the cryoPROS-assisted refinement and particle addition workflow used to improve the resolution of the electron potential map from 7.87 Å to 4.9 Å. Particle subsets obtained from data processing using Relion-4.0 were independently processed through neural network training, auxiliary particle generation, preferential orientation mitigation, non-uniform refinement, particle selection and iterative post-processing steps. Further local refinement, masking and FSC-based validation improved the map quality to 4.9 Å. The sphericity and FSC plots are also displayed.

**Figure 5:**
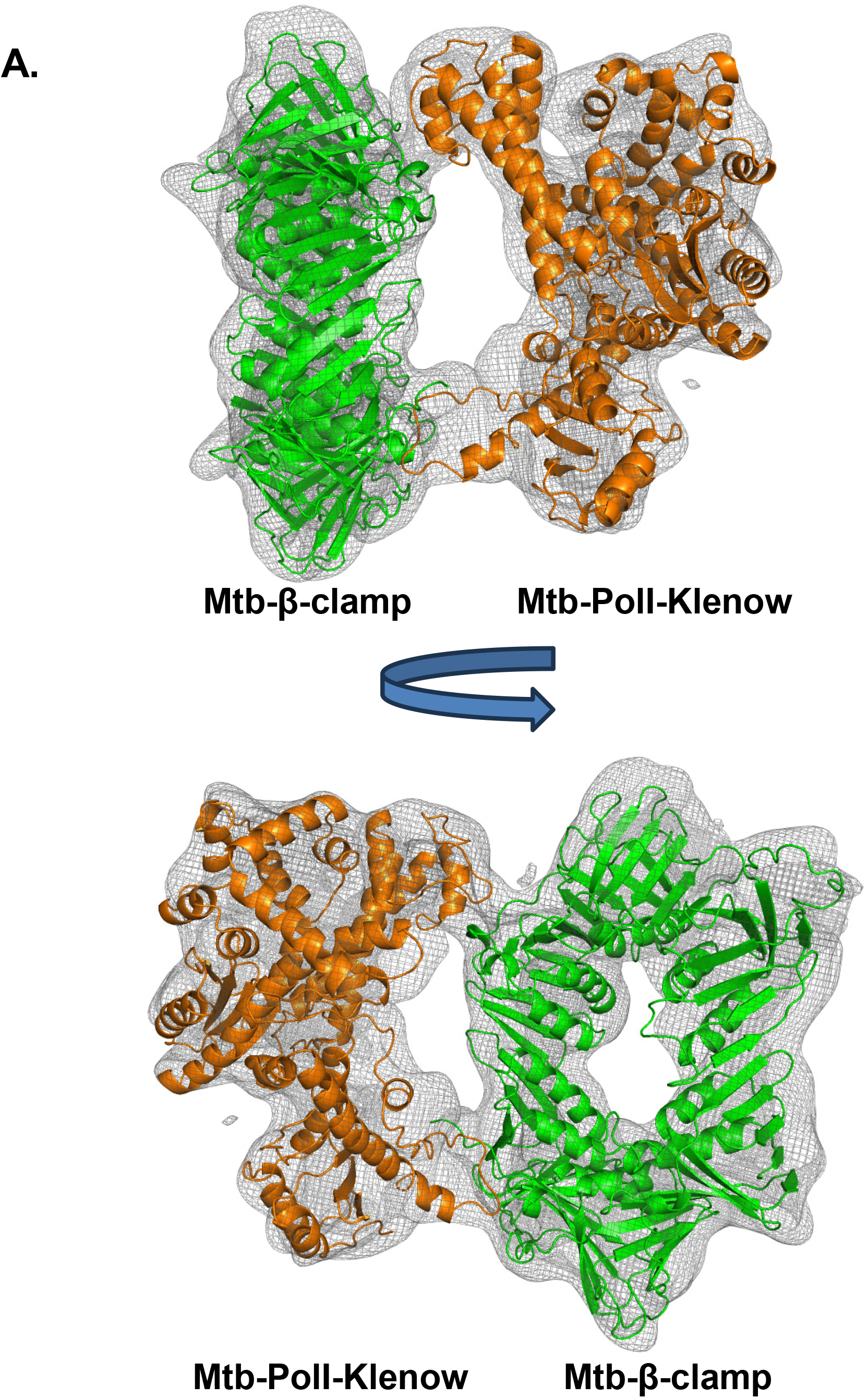
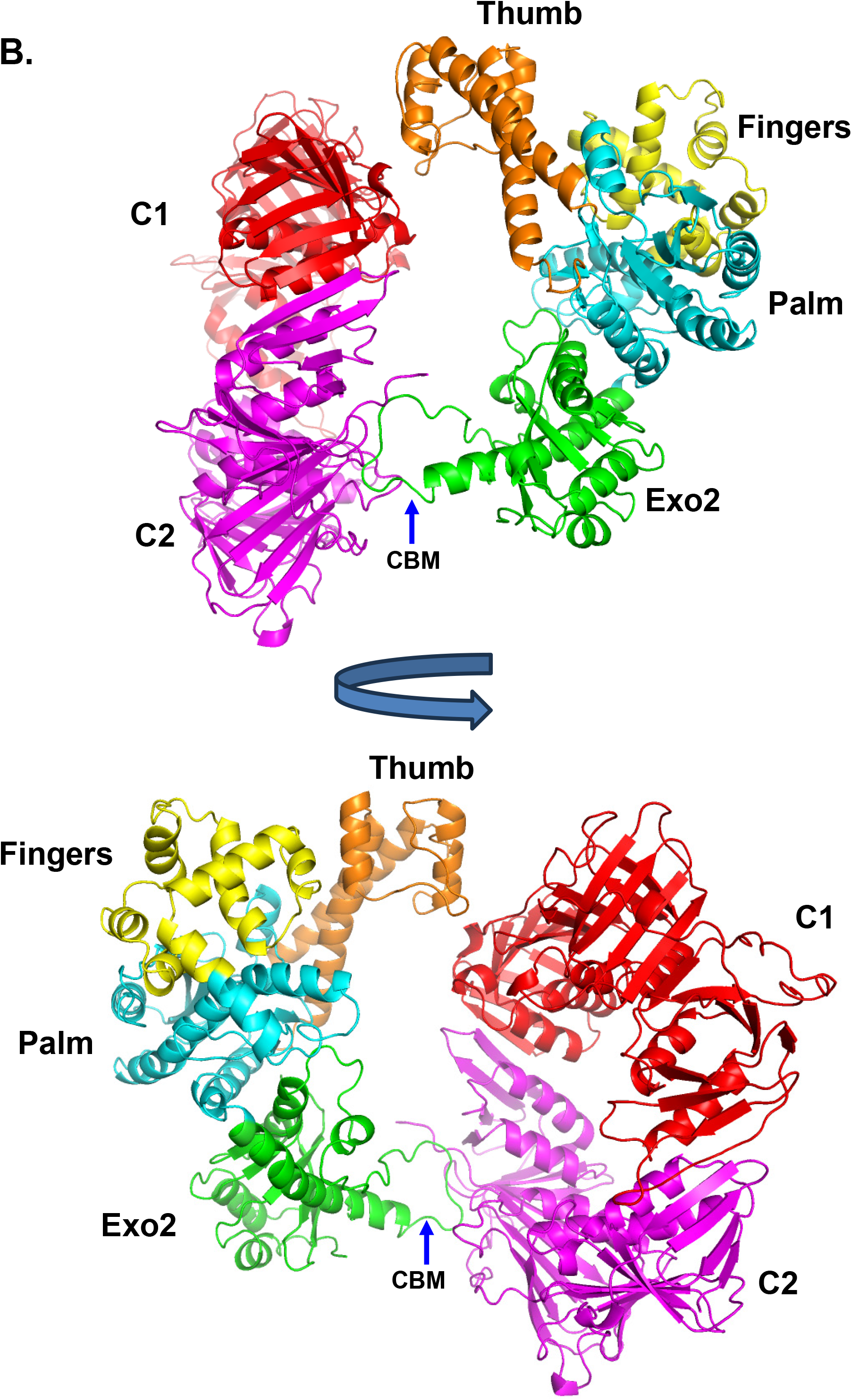
Structure of the Mtb-PolI-Klenow:Mtb-β-clamp complex. (A) The electron potential map, fitted with the structures of Mtb-β-clamp (green) and Mtb-PolI-Klenow (orange), is displayed here in two orientations related by a 90° rotation about the Y-axis. The map is shown in mesh representation, while the protein molecules are shown in cartoon representation. (B) Structure of the Mtb-PolI-Klenow:Mtb-β-clamp complex in cartoon representation is displayed. The proofreading domain and the thumb, palm, and fingers subdomains of Mtb-PolI-Klenow are coloured green, orange, cyan, and yellow, respectively. The monomers C1 and C2 of Mtb-β-clamp are coloured red and magenta, respectively. The blue arrows highlight the location of the <u>C</u>lamp-<u>b</u>inding-<u>m</u>otif (CBM) located between the EXO2 and POL domains.

**Table 1:** Data collection & Validation statistics for the cryoEM structure of the Mtb-PolI-Klenow:Mtb-β-clamp complex.

| **Collection Parameters** |  |
| --- | --- |
| Instrument | Titan Krios |
| Camera | FEI FALCON III (4k x 4k) |
| Recording Mode | Counting |
| Image dimensions (px) | 4096 x 4096 |
| Pixel size ($\text{\AA}/\text{px}$) | 1.07 |
| High Tension (kV) | 300 |
| Energy filter slit width (eV) | No used |
| Cs (mm) | 2.7 |
| Total Dose ($\text{e}^-/\text{\AA}^2$) | 28.81 and 28.35 |
| Exposure time (s) | 60 |
| Frame time (s) | 2.4 |
| Defocus range ($\mu\text{M}$) | -1.8 to -3.5 |
| Processing | Relion and cryoSPARC |
| Images collected | 2991 |
| Images processed | 2945 |
| Particles extracted | 25,98,216 |
| Box size (px) | 256 |
| Consensus particles | 1,39,196 |
| FSC resolution at 0.143 cutoff | 4.9 |
| Sharpening B-factor applied | -426 |
| **Validation Statistics** |  |
| Number of atoms | 10275 |
| Protein residues | 1372 |
| RMS (bonds) | 0.12 |
| RMS (angles) | 0.36 |
| Ramachandran favored | 95% |
| Ramachandran allowed | 5% |
| Ramachandran outliers | 0% |
| Rotamer outliers | 0% |
| Clash score | 14 |
| Q-Score | 0.143 |

The peptide stretch ^456^QQLSLLDDD^464^ from PolI is located in the canonical binding site of one of the monomers (A) of the dimeric Mtb-β-clamp (Fig. 5). The canonical binding site is in domains II (131-262) and III (263-402) of the processivity β-clamp and involves the protein’s C-terminal end. To validate the central role of the ^456^QQLSLLDDD^464^ motif in interaction with the Mtb-β-clamp, site-specific mutations were introduced to replace this motif with ^456^QQASAADDD^464^ in Mtb-PolI-FL and Mtb-PolI-Klenow. The mutant proteins were named Mtb-PolI-FL-mCBM and Mtb-PolI-Klenow-mCBM, and their ability to bind Mtb-β-clamp was assessed. MST experiments showed that Mtb-PolI-Klenow-mCBM and Mtb-PolI-FL-mCBM did not exhibit any measurable interaction with the processivity clamp (Fig. 6A & 6B). Gel filtration analysis showed that, in contrast to the wt proteins, which eluted as a separate peak with lower retention volume, the mixture of Mtb-β-clamp and Mtb-PolI-Klenow-mCBM gave a single broad peak with a retention volume similar to that of individual proteins (Fig. 6C & 6D). Hence, Mtb-PolI-Klenow-mCBM did not form a stable complex with the Mtb-β-clamp dimer. Therefore, the mutations L458A, L460A, and L461A disrupted the complex, and these observations are consistent with the assertion that the ^456^QQLSLLDDD^464^ region represents the primary interaction site on PolI.

**Figure 6:**
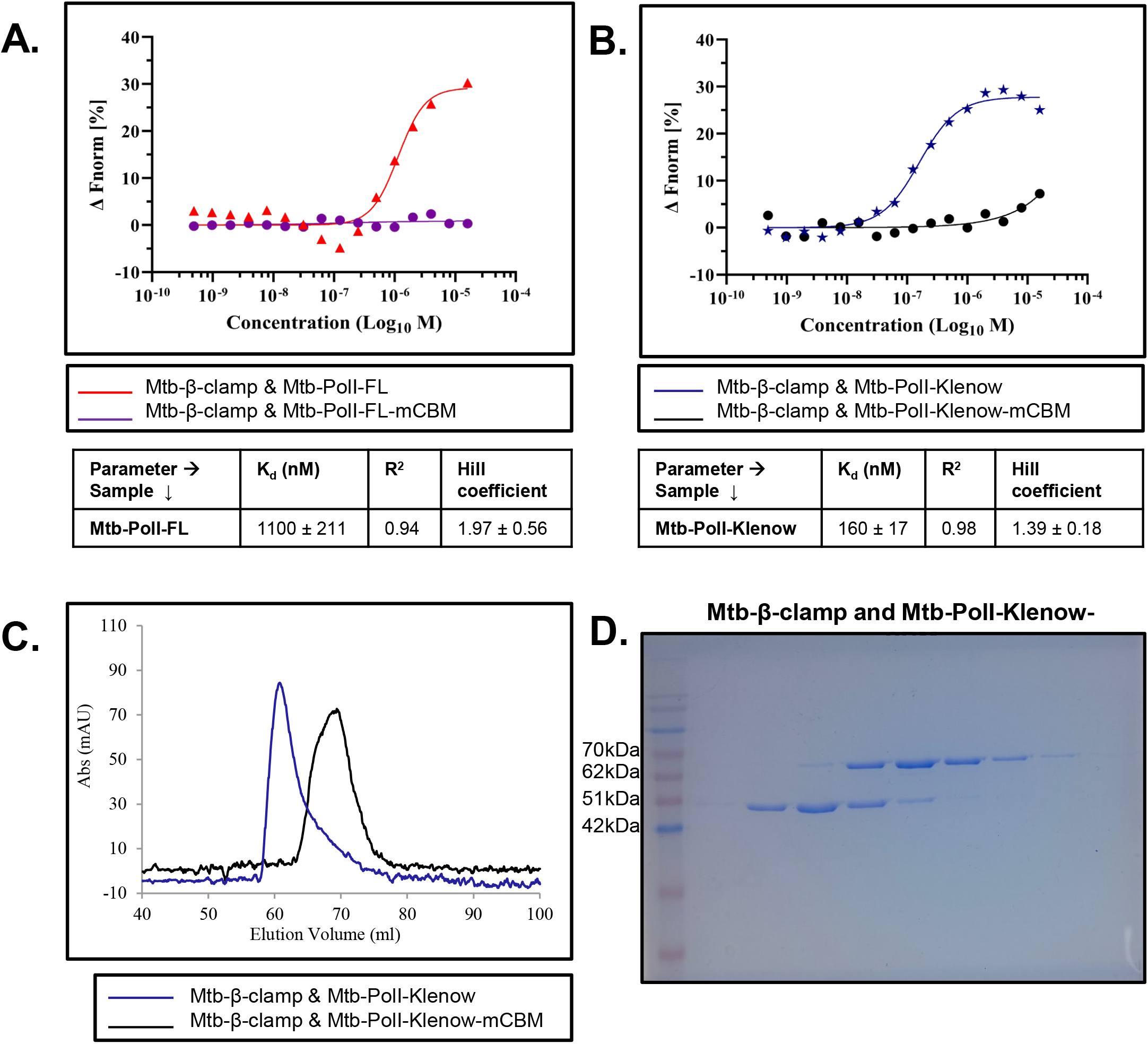
Mutational Analysis of the Mtb-PolI CBM: Binding curves generated using MST data are displayed for the interaction of Mtb-β-clamp with (A)Mtb-PolI-FL (red) and Mtb-PolI-FL-mCBM (purple) and (B) Mtb-PolI-Klenow (blue) and Mtb-PolI-Klenow-mCBM (black). (C) Gel filtration analysis for the mixture of a Mtb-β-clamp and Mtb-PolI-Klenow-mCBM (black). For comparison, the previously obtained profile with Mtb-PolI-Klenow:Mtb-β-clamp complex is also displayed (blue). (D) SDS-PAGE analysis of the eluted fractions for the mixture of Mtb-β-clamp and Mtb-PolI-Klenow-mCBM is displayed.

### Interaction between Mtb-PolI CBM peptide and Mtb-β-clamp

Fluorescence anisotropy studies conducted with 6-FAM-labelled QLSLLDD peptide showed that this peptide binds Mtb-β-clamp with a K_d_ value of 272 nM, which is comparable to the affinity measured between Mtb-PolI-Klenow and Mtb-β-clamp (Fig. 7A). In comparison, the scrambled peptide DLDSLQL did not show any interaction, indicating that binding of the QLSLLDD stretch in Mtb-PolI to Mtb-β-clamp is sequence-specific, further reinforcing its identification as a CBM (Fig. 7B). Interestingly, the shorter peptide (QLSLLDD) shows a stable interaction with Mtb-β-clamp than the longer peptide (QQQQLSLLDDDD), suggesting that the core determinant required for the interaction with Clamp is present within the QLSLLDD stretch (Fig. 7C).

**Figure 7:**
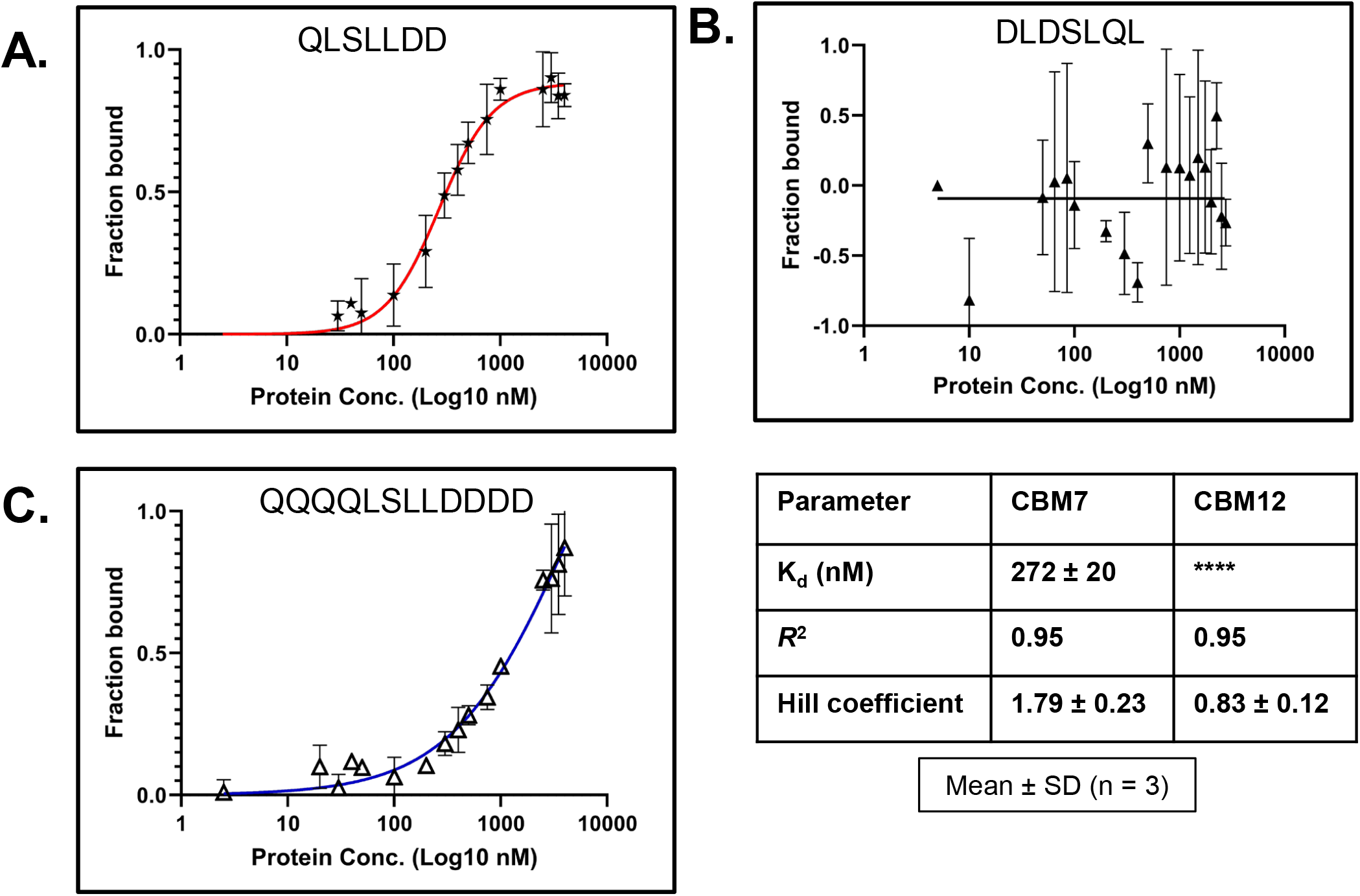
Interaction of Mtb-β-clamp interaction with CBM peptides: The binding curves obtained using fluorescence anisotropy for the interaction between Mtb-β-clamp with CBP7 (red) and CBP12 (blue) peptides, along with the scrambled version CBP7 (black), are displayed, and the interaction parameters are shown in the table.

Crystallisation trials using the hanging drop method were conducted to obtain crystals of Mtb-β-clamp with the unlabeled CBP7 and CBP12 peptides. After extensive optimisation, the best crystals were obtained with CBP12 using a well solution containing 25% PEG 4k, 0.1 M HEPES at pH 7.5, and 0.7 M sodium acetate. The best dataset was collected at the PXIII beamline of the Swiss Light Source to a maximum resolution of 2.9 Å, and the data statistics are provided in Table 2. The structure was solved by Molecular Replacement using 3RB9 as the search model, and the initial maps showed clear density in the canonical binding site. The structure showed the presence of two clamp dimers in the asymmetric unit, and the monomers were named-A, B, C and D. The best density for the CBP12 peptide was observed for the A monomer, and the eight residues of the peptide (AAQLSLLDD) could be built into the density. The Polder map for the peptide bound to the A monomer is displayed at a contour of 4.5^24^ (Fig. 8A).

**Figure 8:**
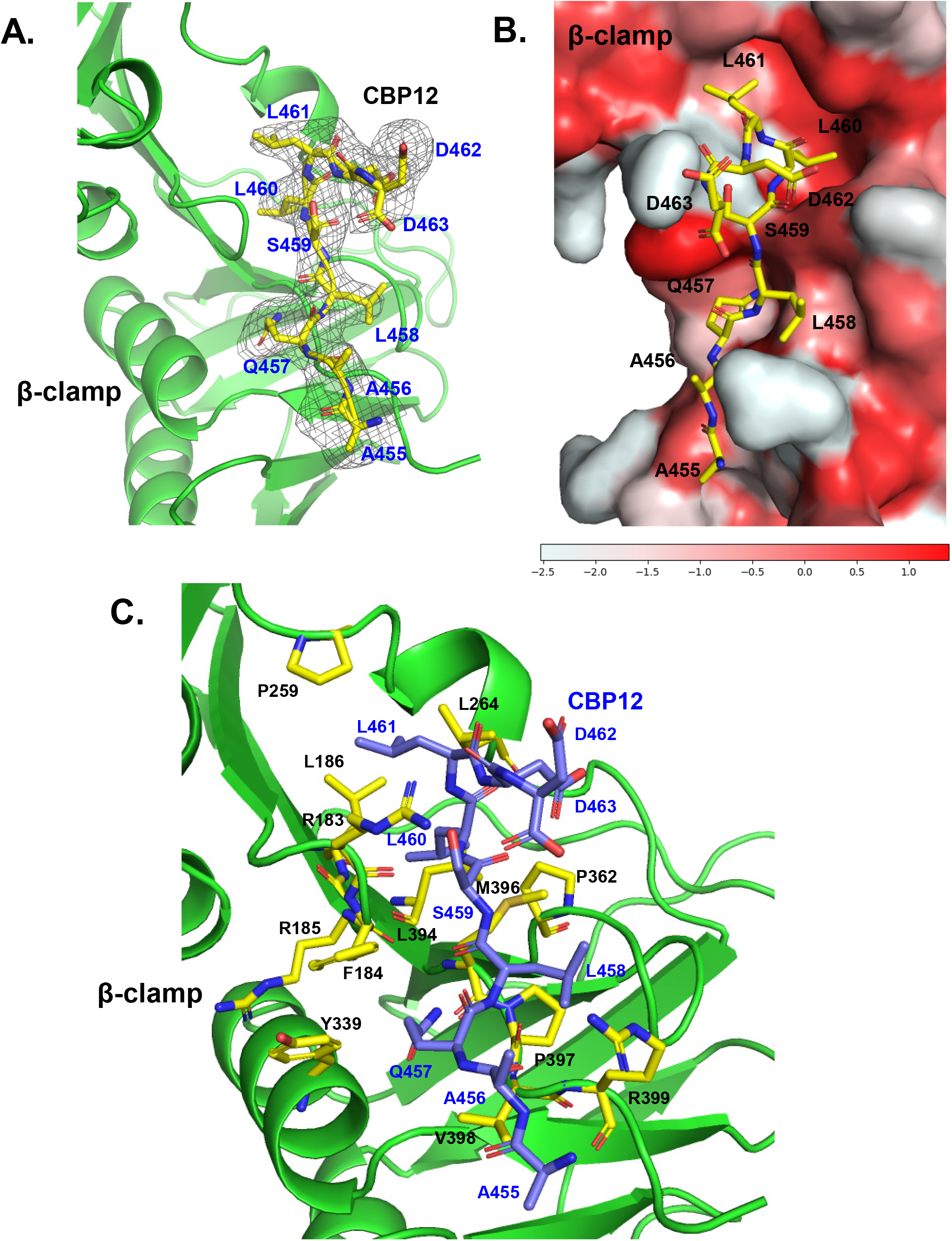
Structure of the Mtb-β-clamp:CBP12 peptide. (A) The C1 monomer Mtb-β-clamp (green) bound to the CBP12 peptide is shown. The polder map for the peptide is displayed as a grey mesh at a 4.5σ contour, and clear electron density was observed for 8 out of the 12 residues of the CBP12 peptide. (B) The surface representation of the canonical binding site on the Mtb-β-clamp is displayed and coloured according to the Eisenberg hydrophobicity scale^76^ (-2.5-hydrophilic to 1.38-hydrophobic). The eight residues of the CBP12 peptide for which electron density was visible are shown in stick representation. (C) The residues of the Mtb-β-clamp that form interactions and the residues of the CBP12 peptide are displayed in stick representation, and coloured by atom with carbon in yellow and blue, respectively. The residues of the peptide are labelled in dark blue, while the residues of the clamp are labelled in black.

**Table 2:**
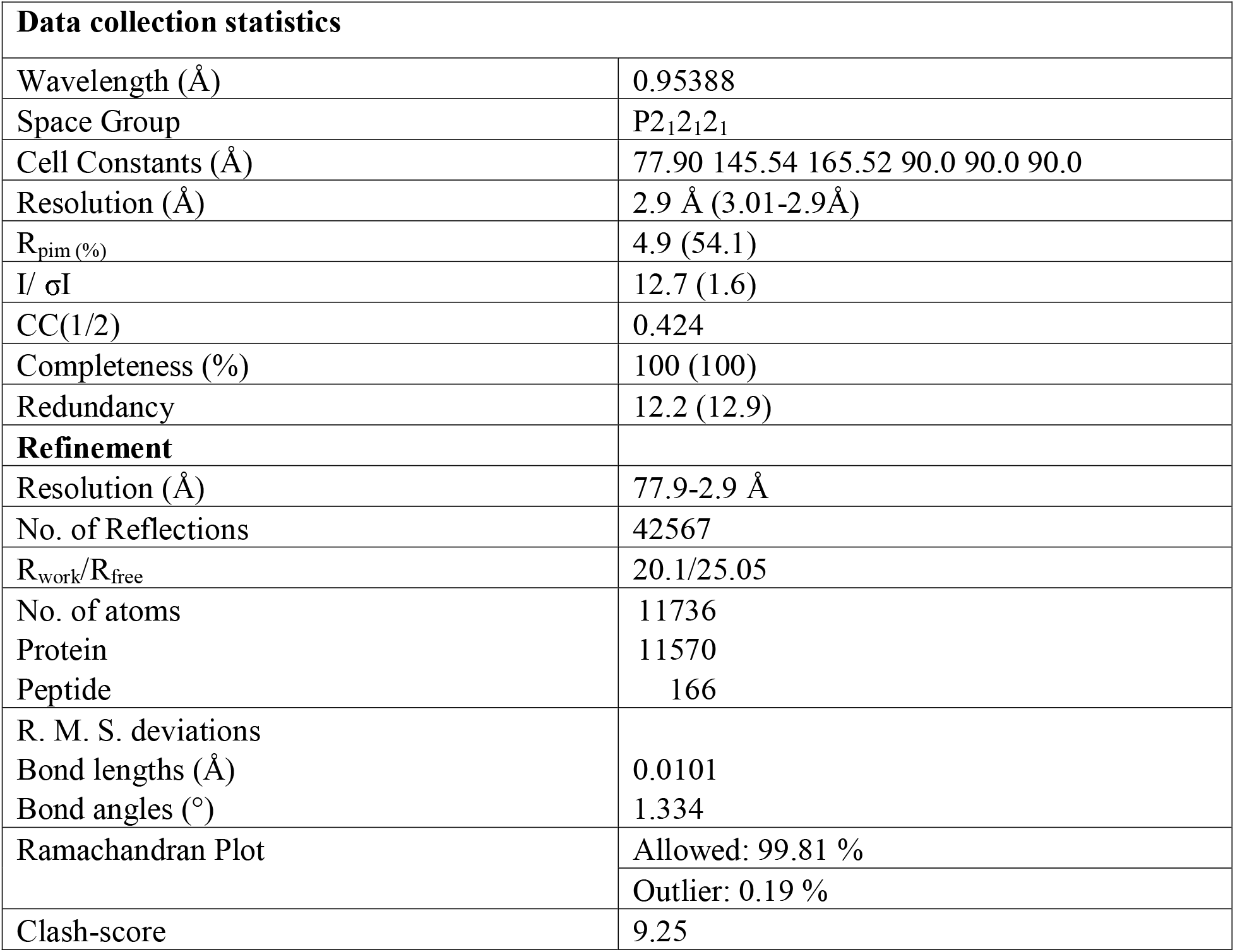
Crystallographic data and refinement statistics for Mtb-β-clamp:CBP12 peptide complex.

| **Data collection statistics** |  |
| --- | --- |
| Wavelength (Å) | 0.95388 |
| Space Group | P2<sub>1</sub>2<sub>1</sub>2<sub>1</sub> |
| Cell Constants (Å) | 77.90 145.54 165.52 90.0 90.0 90.0 |
| Resolution (Å) | 2.9 Å (3.01-2.9Å) |
| R<sub>pim</sub> (%) | 4.9 (54.1) |
| I/ σI | 12.7 (1.6) |
| CC(1/2) | 0.424 |
| Completeness (%) | 100 (100) |
| Redundancy | 12.2 (12.9) |
| **Refinement** |  |
| Resolution (Å) | 77.9-2.9 Å |
| No. of Reflections | 42567 |
| R<sub>work</sub>/R<sub>free</sub> | 20.1/25.05 |
| No. of atoms | 11736 |
| Protein | 11570 |
| Peptide | 166 |
| R. M. S. deviations |  |
| Bond lengths (Å) | 0.0101 |
| Bond angles (°) | 1.334 |
| Ramachandran Plot | Allowed: 99.81 % |
|  | Outlier: 0.19 % |
| Clash-score | 9.25 |

The backbone structure of the Mtb-β-clamp homo-dimers is similar to that of previously deposited structures, and the AB and CD homodimers superimpose onto those deposited in the PDB with accession codes 3P16, 3RB9, 4TR7, 5AGU, 6FVN and 6FVO, with RMSD values in the range of 0.66 to 1.45 Å. Bacterial processivity clamps generally interact with a peptide stretch of the partner protein via contiguous binding sites formed by residues from the clamp monomer’s domain II and the C-terminal domain III. The interaction between residues of the ^457^QLSLLDD^463^ peptide and the Mtb-β-clamp binding site is largely hydrophobic, augmented by a few polar interactions (Fig. 8B and Supplementary Movie S4). For Mtb-PolI-Klenow and Mtb-PolI-FL, the mutations L458A, L460A, and L461A abolished the enzymes’ ability to bind the clamp. The corresponding three leucine residues in the CBP12 peptide are present in cavities lined by apolar residues of the clamp, such as F184, M396, L394, L186 and P259 (Fig. 8C). The side chain of the Q457 residue forms hydrogen bonds with the side chain of N336 and the backbone carbonyl of M396. There are hydrogen bonds formed between the backbone atoms of Q457, L458, and L460 from the peptide and the Mtb-β-clamp residues R399, P397 and R183, respectively. The backbone atoms of D462 and D463 form hydrogen bonds with the side chain of R183.

The CBM peptide-binding site analysis, along with the crystal structure of the Mtb-β-clamp:CBP12 complex, revealed that the interaction between Mtb-PolI and the processivity clamp is sequence-specific. The stretch ^457^QLSLLDD^463^ of Mtb-PolI, therefore, represents a *bona fide* Clamp-binding motif.

### Phylogenetic analysis of PolI sequences

A pairwise alignment of the PolI sequences from Mtb and *E. coli* shows that the QLSLLD motif is missing in the latter, and consequently, the interaction with the processivity clamp is weak. The available phylogenetic analysis, prepared using 151 representative bacterial species, was utilised (Fig. 9A), to select 25 organisms distributed across the clade for sequence analysis to detect the presence of the QLSLLDD motif^25^. This analysis showed that the PolI sequence from members of the Mycobacteriaceae and Corynebacteriaceae families possessed sequences homologous to the QLSLLDD motif between the EXO2 and POL domains (Fig. 9B). According to NCBI taxonomy (Taxon ID 85007), these two families are classified in the Mycobacteriales order.

**Figure 9:**
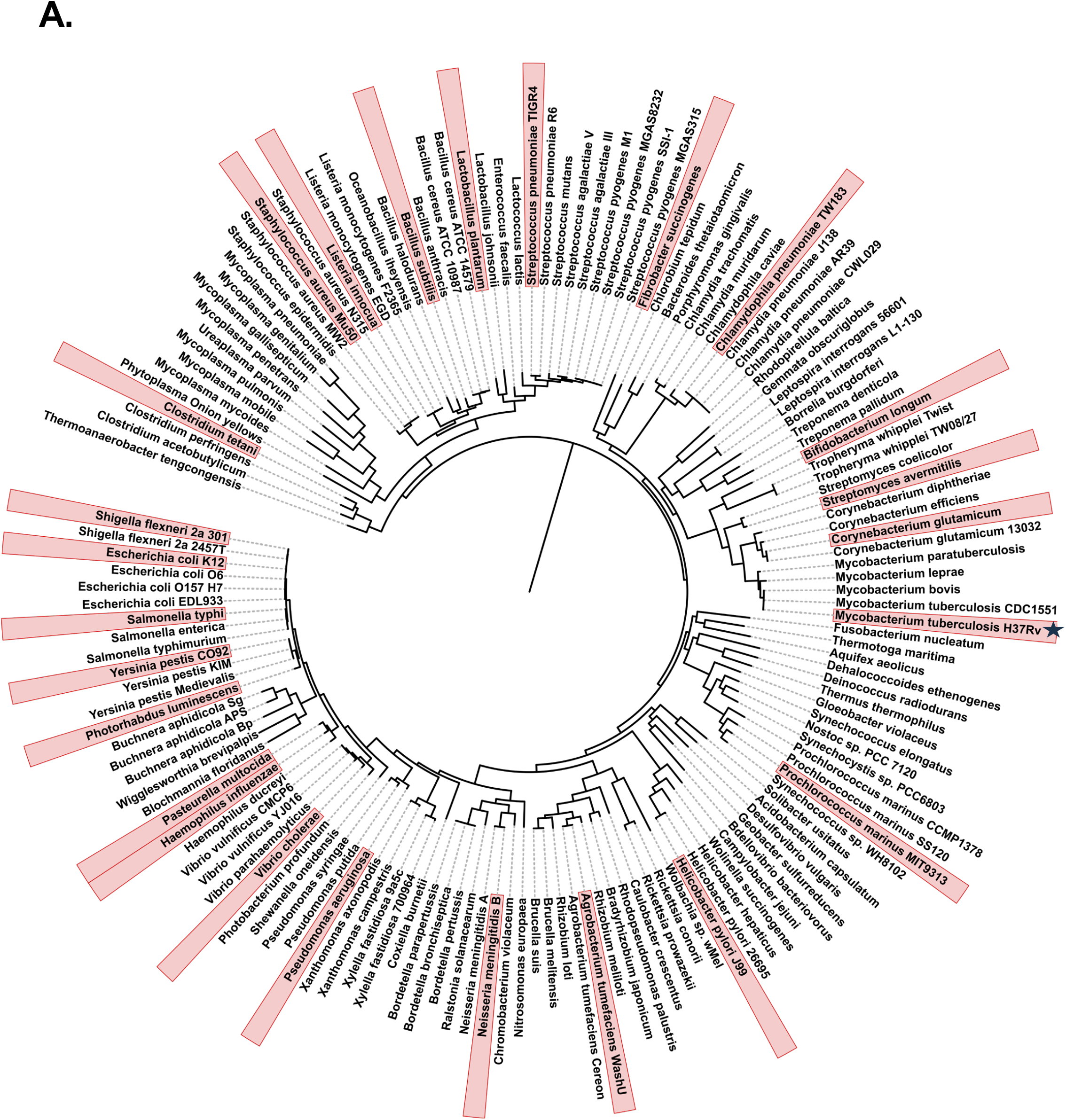
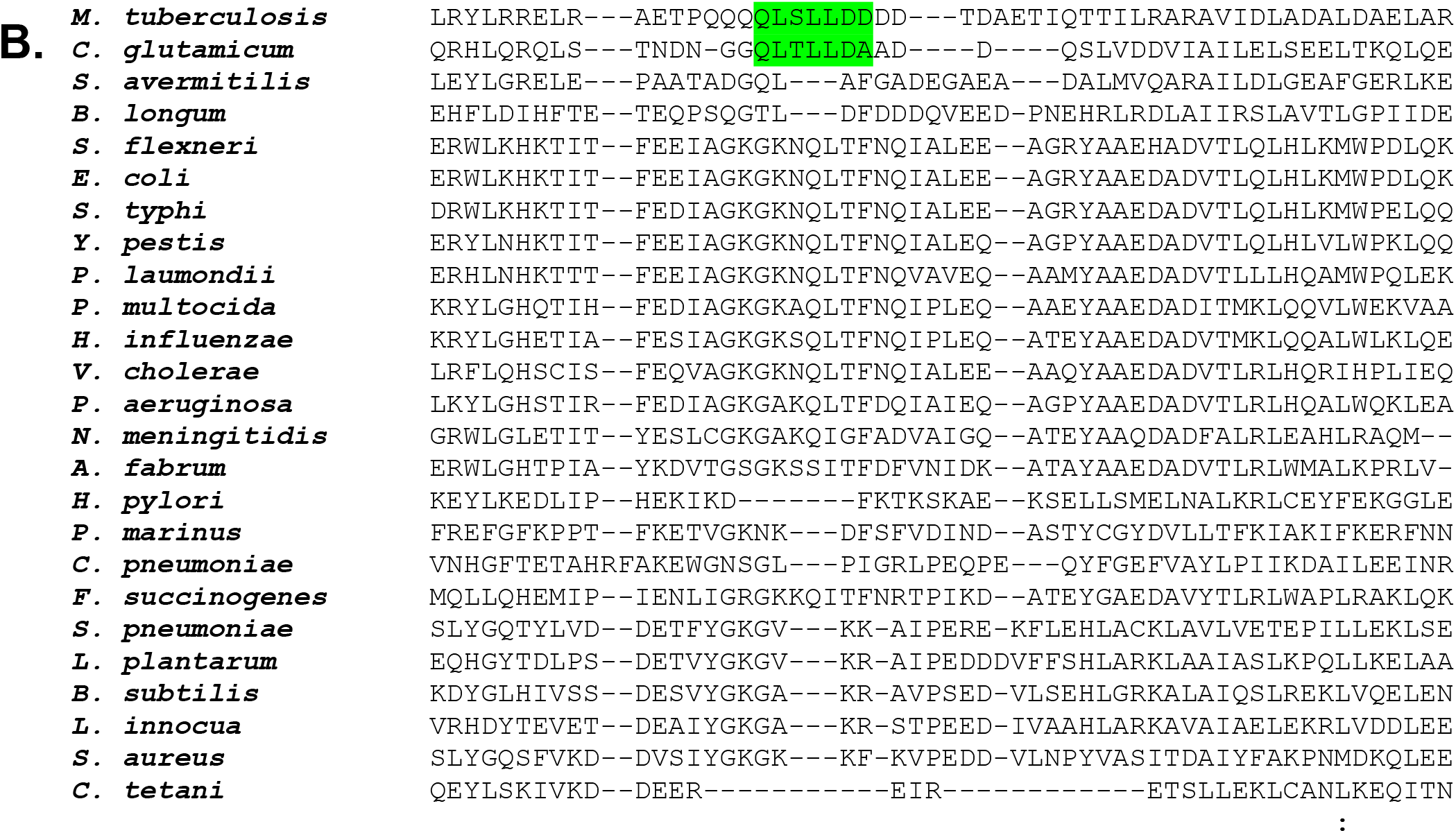
Phylogenetic analysis and Multiple sequence alignment. (A) The phylogenetic tree generated from 151 representative bacterial species is shown, with the selected species for further analysis highlighted and Mtb marked with a star. (B) The multiple sequence alignment of the polypeptide stretch encompassing the CBM in Mtb is displayed for the selected 25 species.

In addition to these two families, the order Mycobacteriales includes the following families: Dietziaceae, Gordoniaceae, Hoyosellaceae, Lawsonellaceae, Nocardiaceae, Segniliparaceae, Speluncibacteraceae, and Tsukamurellaceae. The PolI and β-clamp sequences from representative members of all these families were aligned. The sequences aligned included those of *Mycobacterium tuberculosis* (Mycobacteriaceae), *Corynebacterium ammoniagenes* (Corynebacteriaceae), *Dietzia aurantiaca* (Dietziaceae), *Gordonia desulfuricans* (Gordoniaceae), *Hoyosella rhizosphaerae* (Hoyosellaceae), *Lawsonella clevelandensis* (Lawsonellaceae), *Nocardia halotolerans* (Nocardiaceae), *Segniliparus rugosus* (Segniliparaceae), *Tomitella fengzijianii* (Tomitellaceae), *Tsukamurella sputi* (Tsukamurellaceae), and *Speluncibacter jeojiensis* (Speluncibacteraceae). It was observed that in PolI from all these organisms, a sequence similar to the Mtb-PolI CBM was present at a similar location between the EXO2 and POL domains (Fig. 10A), and the consensus sequence is - QLSLLDD (Fig. 10B). Also, residues of Mtb-β-clamp involved in interaction with PolI CBM are conserved among the selected families (Supplementary Fig. S4). Overall, the phylogenetic analysis and multiple sequence alignments of interaction interface residues of PolI and β-clamp suggest that the strong interaction observed for proteins from Mtb is evolutionarily conserved across bacteria from different families within the Mycobacteriales order in the Actinomycetes class.

**Figure 10:**
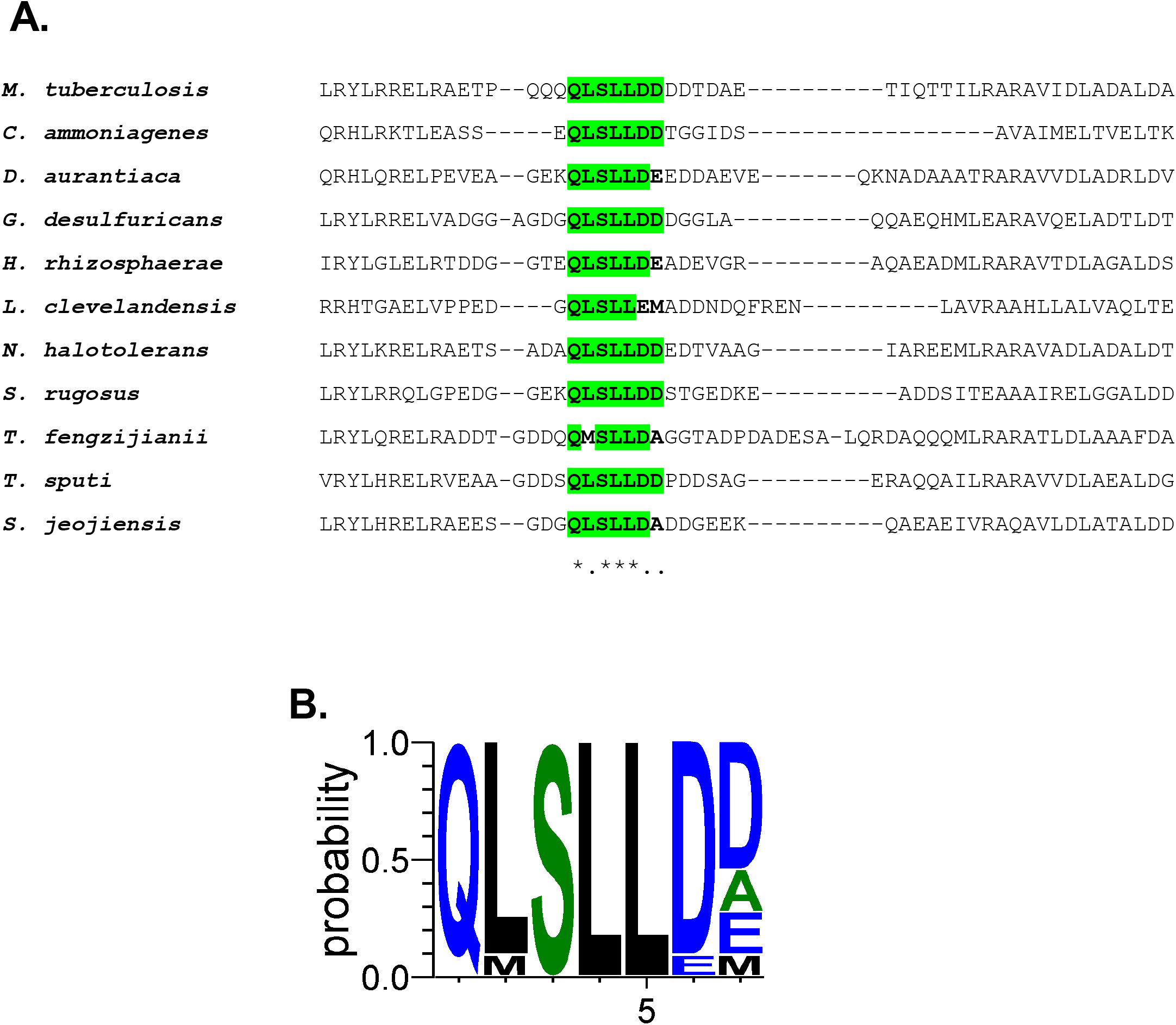
Conservation analysis of CBM in PolI from members of the order Mycobacteriales: (A) Multiple sequence alignment (MSA) of the region encompassing the CBM in PolI sequences from representative members of the Mycobacteriales order is displayed. The conserved residues in the CBM of PolI are highlighted in green. (B) Based on the MSA, a graphical summary of the position-specific conservation and residue frequencies for the CBM of PolI is displayed.

## Discussion

Our study shows that PolI interacts with β-clamp through the Klenow fragment region and forms both stable and transient interaction interfaces. The resolution of the electron potential map generated from the available cryoEM data could be considerably improved with cryoPROS, which employs a self-supervised deep generative model to reduce preferred-orientation–driven misalignment. Also, continuous heterogeneity analysis clearly showed that the thumb domain moves, and hence the interaction between the thumb residues and the non-canonical site on the clamp is secondary. The use of recently released tools, such as 3DFlex and cryoPROS, provided dynamic information and improved the resolution of the final electron potential map, respectively, thereby providing more in-depth information from the available cryoEM data. The stable, sequence-specific interaction between Mtb-PolI and Mtb-β-clamp involves the CBM located between the EXO2 and POL domains, and this CBM is evolutionarily conserved within PolI from members of the Mycobacteriales order.

The cryoEM structure shows that only one molecule of PolI binds to the clamp dimer, whereas the crystal structure shows that all four monomers of Mtb-β-clamp present in the asymmetric unit are bound with peptide. It appears that dynamic interactions with the thumb domain occlude the second canonical binding site on the clamp dimer. Also, there is a difference in the affinities of CBM peptide, Mtb-PolI-Klenow and Mtb-PolI-FL for the Mtb-β-clamp. The comparatively strong interaction with the Klenow fragment may arise from simultaneous contributions from two interaction interfaces, whereas the CBM represent only a single interaction determinant and therefore displays reduced binding strength. In contrast, the weaker interaction observed for full-length PolI suggests that the EXO1 domain may negatively modulate the accessibility or dynamics of Mtb-PolI-FL:Mtb-β-clamp via the CBM peptide, thereby playing a regulatory role.

Among prokaryotes, the β-clamp generally displays a conserved architecture and interacts with the CBM of partner proteins via the canonical binding site on each monomer. The α-subunit of the DNA polymerase III holoenzyme interacts with the β-subunit through two peptides defined by ^920^QADMF^924^ and ^1154^QVELEFD^1160^. The B-family polymerase, DNA polymerase II, binds the clamp via the ^779^QLGLF^783^ motif^23,26,18,27^. The Y-family dPols-DNA polymerase IV and V-interact with the β-subunit through the ^346^QLVLGL^351^ and ^357^QLNLF^361^ motifs, respectively^28–30^. The general consensus of the clamp-binding motif (CBMs) is Qφχ [L/M][F/L] or Qφ χ [L/M] χ [F/L], where φ is an aliphatic residue and χ any residue^19^. The CBM identified in Mtb-PolI does not fully match the consensus sequence, although the core “QLSLL” segment is present. The Mtb-PolI CBM sequences is closer to the consensus sequence QL[SD]LF previously identified by Dalrymple et al^23^. In Mtb-PolI, the QLSLLD motif is part of the sequence ^454^QQQQLSLLDDDD^465^, and the low-complexity stretches before and after the CBM may prevent the formation of secondary structure in this region, thereby ensuring easy access of the CBM to the binding site on the clamp for rapid and stable engagement. Among mycobacterial dPols, the QLSLLD motif is unique to PolI and is not present in DNAE1 (Rv1547), DNAE2 (Rv3370), DinB1 (Rv1537), DinB2 (Rv3056) and PolX (Rv3856c).

The stable interaction between PolI and the β-clamp is a unique feature of genome replication and repair in Mycobacteria, distinct from that in *E. coli*. Although there are similarities in the general strategy for genome replication and repair across the two organisms, there are multiple points of divergence, which are likely the result of differences in their environments and replication rates^31,32^. Deletion of the *polA* gene is known to increase sensitivity to DNA-damaging agents in *M. smegmatis* ^8^. Since Mtb is continuously exposed to DNA-damaging agents within the host cell, the function of PolI and, by extension, its interaction with DnaN may be crucial for Mtb survival in macrophages^32^. The “toolbelt model” describes how different dPols can engage simultaneously with the processivity clamp at the replication fork to rapidly switch between high-fidelity replication and low-fidelity translesion DNA synthesis^33^. The ability of PolI to interact strongly with the β-clamp will allow this enzyme to participate in the toolbelt model, facilitating rapid switching between different functions associated with dPols, such as DNA synthesis and processing of RNA-DNA hybrids. The NucS protein, which is conserved in the Mycobacteriales order, is involved in the non-canonical mismatch repair pathway of mycobacteria^34^, and this function is mediated through interaction with the β-clamp^35,36^. NucS cleaves mismatches in DNA, leading to a short patch that must be resected and resynthesized. Due to its ability to interact with the β-clamp, PolI in members of the order Mycobacteriales may participate in resection and synthesis during NucS-associated genome maintenance pathways. Overall, the stable interaction between PolI and the β-clamp, mediated by the unique CBM, may enable PolI to play a greater role in genome repair in the order Mycobacteriales.

The bacterial processivity clamp is an essential protein that exhibits oligomeric assembly and a canonical binding site distinct from that of the eukaryotic homolog, proliferating cell nuclear antigen^37,38^. For these reasons, the β-clamp represents an attractive drug target, and a number of lead molecules have been identified that bind to block the canonical binding site on the processivity clamp^19,27,39–41^. The Mycobacteriales order includes a number of pathogens, including Mtb, for which the emergence of new drug-resistant strains is of global concern, as well as other Mycobacteria such as *M. leprae*, *M. abscessus*, and *M. ulcerans* (Supplementary Fig. S5). Other emerging pathogens classified in the order Mycobacteriales include *Nocardia brasiliensis* and *Gordonia bronchialis*, which can cause severe disease in immunocompromised patients^42,43^. The conserved new peptide motif described in our study can aid the development of peptidomimetic molecules that bind the processivity clamp of pathogenic members of the order Mycobacteriales and perturb normal replication and repair. Our studies may therefore facilitate the structure-based drug discovery efforts targeting DNA replication and repair pathways in multidrug-resistant mycobacteria.

## Materials and methods

### Cloning, expression and purification of wt and mutant proteins

The gene and gene segments corresponding to PolI (Rv1629) and the corresponding Klenow fragment, respectively, from *Mycobacterium tuberculosis* (Mtb) H37Ra were amplified from genomic DNA and cloned into the pGEX-6P-1 vector using the EcoRI/XhoI sites. Site-directed mutagenesis was used to generate a mutant version of Mtb-PolI-FL that is deficient in EXO1 activity, and this clone was subsequently used to generate a construct corresponding to the EXO1 domain. The gene for PolI from *E. coli* was amplified from genomic DNA (DH5-α cells) and cloned into the pGEX-6P-1 vector between the EcoRI/NotI sites. The *dnan* gene from Mtb and *E. coli* was cloned into a modified pET-28b vector between BamHI/NotI. The gene construct corresponding to the Klenow fragment of PolI from *E. coli* used in our studies was already available in the lab ^44^. Site-specific mutations used in this study were generated with the QuikChange Lightning kit (Agilent Technologies) according to the manufacturer’s instructions. All the clones and mutations were confirmed with Sanger sequencing.

The cloned constructs for Mtb-PolI-FL, Ec-PolI-FL, Mtb-PolI-Klenow, Ec-PolI-Klenow, Mtb-β-clamp, Ec-β-clamp and Mtb*-*PolI*-*EXO1 exhibited optimal expression in the C41 (DE3) strain of *E. coli*. For each construct, cells were harvested from 5 litres of culture, induced with IPTG, and incubated overnight at 18 with shaking. The cells were lysed by sonication, the lysate was clarified by centrifugation, and the filtered supernatant was used for affinity chromatography. For the proteins expressed using the pGEX-6P-1 vector, GST-sepharose was used; for the other proteins, charged Nickel-NTA resin was used, following standard protocols^12,45^. The tags were cleaved with PreScission protease, and the proteins were further purified by size-exclusion chromatography on a 16/600 Superdex 200 column (Cytiva Inc.). The fractions were analysed using SDS-PAGE, and those corresponding to pure protein were pooled, concentrated, flash-frozen, and stored at -80 °C. The mutants of Mtb-PolI-Klenow and Mtb-PolI-FL were purified using a protocol similar to that for wild-type.

### MicroScale Thermophoresis (MST)

MST was performed to assess the interactions between Mtb-β-clamp and Mtb-PolI-EXO1, Mtb-PolI-FL, and Mtb-PolI-Klenow, as well as mutant versions of the latter two proteins. All protein samples used for the MST study were buffer-exchanged into the MST buffer (25 mM HEPES, pH 7.5 at 4 °C, 50 mM NaCl, 5% Glycerol, 2 mM DTT, and 0.05% Tween 20). Mtb-β-clamp protein was labelled with RED-tris-NTA dye as per the manufacturer’s protocol. Labelled Mtb-β-clamp was incubated with varying concentrations (from 16000 nM to 0.48 nM) of different constructs of Mtb-PolI, and the fluorescence-based thermoporetic signal was measured using the Monolith NT.115 instrument (NanoTemper GmbH). The percentage change in normalised fluorescence (ΔFnorm in %) was plotted on the Y-axis, and the log_10_ (Concentration in nM) of the target protein was plotted on the X-axis. The data were analysed in GraphPad Prism (version 10.3.1) using nonlinear regression specific binding with the Hill slope equation to determine the binding affinity (K_d_). A similar protocol was used to quantify the binding affinities of Ec-PolI-FL and Ec-PolI-Klenow for Ec-β-clamp.

### Analytical gel filtration chromatography

The 16/600 Superdex 200 column (Cytiva Inc) was equilibrated with the buffer-25 mM HEPES, pH 7.5, at 4 °C, 50 mM NaCl, 5% Glycerol and 2 mM DTT. 400 µg of the β-clamp from Mtb or *E. coli* was mixed with a molar equivalent of the corresponding PolI-Klenow or PolI-FL proteins. The individual proteins and the incubated mixtures were injected into the column, and the chromatography profiles were recorded. The eluted fractions were run on 12% SDS-PAGE to determine their composition.

### Dynamic light scattering (DLS) analysis

The purified Mtb-β-clamp, Mtb-PolI-Klenow and Mtb-PolI-Klenow:Mtb-β-clamp complex were concentrated to 4 mg/ml. The analytical buffer and the protein solutions were centrifuged at 20,000 g for 45 min at 4 immediately before measurement. The data were collected using a Zetasizer Nano ZS90 (Malvern Instruments Ltd), and all measurements were carried out in a low-volume Quartz cuvette (Malvern Instruments Ltd) at room temperature. The data was processed and analysed using the Zetasizer software associated with the instrument to calculate hydrodynamic diameter (D_H_) values using the Stokes–Einstein equation.

### Cryo-Electron Microscopy data collection and processing

The purified Mtb-PolI-Klenow:Mtb-β-clamp complex (2 mg/ml) was applied to glow-discharged Quantifoil® R 1.2/1.3 Au 300-mesh grids, followed by plunge-freezing using a Vitrobot Mark IV. Two datasets, consisting of 1256 and 1735 movies, were collected at the National Electron Cryo-Microscopy Facility, Bangalore Life Science Cluster (BLiSc), Bangalore, using a 300 kV Titan Krios microscope with a Falcon3 detector.

Initial cryo-EM data were processed using cryoSPARC v4.4.1 ^46^, installed on the BRAHM HPC of the Indian Biological Data Centre (IBDC). Briefly, movies were imported, motion corrected, and CTF estimation was carried out ^47–49^. A total of 138 micrographs were discarded as outliers, and Blob Picking was performed using a particle diameter range of 100– 170 Ǻ, followed by particle extraction with a box size of 256 pixels. After multiple rounds of 2D classification and class selection, a total of 18 classes with 355,473 particles from dataset 1 and 19 classes with 393,988 particles from dataset 2 were further used for *ab initio* reconstruction ^46^. In datasets 1 and 2, only single classes have both proteins sharing 41.3% and 39.4% of particles, respectively. The particles corresponding to the complex from both datasets were combined, and Non-Uniform Refinement (NU-Refine) was performed ^50^. The resulting volume map was further processed using sharpening, volume analysis, and FSC-based validation tools (FSC)^51^. Structural docking was performed with Mtb-β-clamp crystal structure (3RB9) and Mtb-PolI-Klenow models generated using Swiss-Model, using the structure of PolI from *M. smegmatis* as a template ( 6VDE)^52^.

For continuous heterogeneity analysis of the cryoEM data, the 3DFlex suite was used^53^. The particles were subjected to data preparation, mesh generation, and a segmentation mask was generated using Segger v2.5.3 in Chimera^54,55^. Subsequently, training, reconstruction, and trajectory generation were performed to identify conformational variability and domain movements within the Mtb-PolI-Klenow:Mtb-β-clamp complex.

To improve the resolution, cryoPROS^56^ was used after the initial cryoEM reconstruction. For this analysis, the datasets were initially processed in Relion-4.0^57^, and the two datasets, comprising 1256 and 1735 movies, were processed independently as optics groups 1 and 2. Movies were imported, motion corrected, and subjected to CTF estimation using CTFFIND-4.1^58,59^. A total of 46 micrographs were discarded based on the quality of the Thon rings. Particle picking was performed using Laplacian of Gaussian (LoG) based auto-picking, followed by extraction with a box size of 256 pixels. After multiple rounds of 2D classification and class selection, a total of 25 classes with 4,95,955 particles from optics 1 and 16 classes with 5,48,802 particles from optics 2 were obtained.

The previously reconstructed 7.87 Å cryoSPARC map was used as an initial model for 3D-classification with four classes^60^. A complex containing class from each dataset containing 1,09,660 and 1,72,106 particles, respectively, was selected for further processing using cryoPROS. Selected particles were imported into cryoSPARC for independent non-uniform refinement using the 7.87 Å map as a reference^50^. Density maps generated from docked models were converted to a low-resolution map in Relion-4.0 and used, together with raw particles, for cryoPROS neural network training, auxiliary particle generation, and iterative refinement. Successive rounds of particle selection, 2D classification, homogenous reconstruction, and retraining yielded a final particle subset of 85,679 and 53,517 particles from optics group 1 and 2, respectively. The final locally refined map, prepared using both sets of particles, was post-processed with EMReady2^61^, followed by sharpening, volume analysis, and FSC-based validation to generate the final gold-standard FSC curve and half-maps. The sphericity of the cryoEM map was analysed using 3DFSC^62^.

The Mtb-β-clamp crystal structure (3RB9) and the homology model of Mtb-PolI-Klenow were docked into the cryoPROS-assisted improved map using ChimeraX^63^. Initial model fitting was done by Coot^64^, segment-specific refinement by ISOLDE^65^ and finally, real-space refinement was performed using Phenix^66^.

### Interaction of CBM peptide from Mtb-PolI with Mtb-β-clamp

Synthetic 7-mer and 12-mer peptides corresponding to the CBM from Mtb-PolI - named CBP7 and CBP12, respectively and the scrambled version of CBP7 were procured, each labelled with 6-FAM. 20 nM of each peptide was incubated with varying concentrations (5.5 µM to 5 nM) of Mtb-β-clamp in a 96-well plate. Following incubation at room temperature for 10 mins, anisotropy measurements were made using a SpectraMax i3x (Molecular Devices) with excitation at 490 nm and emission at 520 nm at 25. Using GraphPad Prism v10.3.1, the graph of Fraction bound versus log_10_(protein concentration in nM) was plotted, and the K_d_ value and standard error of mean were measured in a logistic equation in Hill’s plot fit.

### Crystallization, Data Collection and Structure Determination

200 µM of Mtb-β-clamp was incubated with 1000 µM of unlabeled CBP7 or CBP12 for 30 min on ice. After extensive screening and optimisation, the best crystals were obtained with CBP12 in 21-24% PEG 4000, 0.4-0.8 M sodium acetate and 0.1 M HEPES, pH 7.5. The plate-like crystals were cryo-protected by soaking in mother liquor containing 5%, 10%, 15%, 20%, 25%, and 30% sucrose for 15 sec, followed by flash-freezing in liquid nitrogen.

For the crystals of the Mtb-β-clamp:CBP12 complex, initial crystal screening was conducted at the home source located at the RCB, and at the BL-21 beamline of the Indus-II synchrotron (RRCAT, Indore, India)^67^. The final dataset was collected at the PXIII beamline of the Swiss Light Source (SLS) to a maximal resolution of 2.9 Å^68^.The data were processed using iMOSFLM^69^, followed by merging and scaling using AIMLESS^70^. The structure was determined by Molecular Replacement with PHASER^71^, using the available Mtb-β-clamp structure (3RB9) as the search model. There were two clamp dimer molecules in the asymmetric unit, and the electron density map was calculated after one round of refinement with PHENIX^66^. The map showed clear density for the peptide in all four clamp molecules of the two dimers. The peptide was built *de novo* using COOT^64^ and multiple rounds of refinement were carried out until the R-factors converged. The interactions between the peptide and clamp were analysed using the CONTACT program in CCP4i^72^. The figures were prepared using either PyMol (Schrodinger Corp.) or ChimeraX, and all the movies were prepared using either Chimera^54^ or ChimeraX^63^.

### Sequence conservation analysis

25 representative species from the bacterial clade were selected based on phylogenetic diversity using the phylogenetic tree available in Interactive Tree Of Life (iTOL)^25^. PolI sequences from the selected organisms were retrieved from the protein database of NCBI^73^ and aligned using Clustal Omega^74^. PolI and β-clamp sequences of representative species from the order Mycobacteriales were retrieved and aligned in a similar manner. The conserved CMB motif identified from the structural and biochemical studies was highlighted in the alignment and further visualised using WebLogo3^75^. Based on the alignment of the β-clamp sequences, the conservation of residues involved in interaction with the CBM of PolI was analysed.

## Acknowledgements

We thank Dr Vinothkumar K.R. from the National Centre for Biological Sciences (NCBS) and Dr Sucharita Bose from the National Electron Cryo-Microscopy facility at the Institute for Stem Cell Science and Regenerative Medicine (inStem) for help in sample preparation and data collection. The National Electron CryoEM facility is supported by the Department of Biotechnology (DBT) B-Life grant DBT/PR12422/MED/31/287/2014. We acknowledge the BRAHM: High-Performance Computational facility of the Indian Biological Data Centre (IBDC), Regional Centre for Biotechnology, Faridabad, INDIA (DBT Grant no. BT/TCB/IBDC/2019), for processing CryoElectron Microscopy data. We thank the X-ray diffraction facility at the Regional Centre for Biotechnology and Dr Shaminder Singh for access to the instrument and technical support. DTN thanks Dr Vincent Olieric (PXIII beamline, Swiss Light Source) for help with X-ray diffraction data collection. The research was supported by the Anusandhan National Research Foundation, Govt. of India, through Grant no. CRG/2022/002638, and by an intramural grant no. C0901 from the Regional Centre for Biotechnology (RCB). We acknowledge the contribution of the Genomics, Molecular Interactions & Electron Facilities of the Advanced Technology Platform Centre (ATPC), which is managed by the Regional Centre for Biotechnology (RCB), and funded by the Department of Biotechnology, Govt. of India (Grant No. BT.MED-II/ATPC/BSC/01/2010). DS was a recipient of a PhD Research Fellowship from the Council of Scientific & Industrial Research, Govt. of India.

## Data Availability

For the Mtb-PolI-Klenow:Mtb-β-clamp complex, the cryo-EM maps have been deposited in the EMDB, and the atomic coordinates have been deposited in the wwPDB with accession codes: EMD-80864 and 26TS, respectively. The Structure factors and the refined coordinates for the Mtb-β-clamp:CBP12 complex have been deposited in the wwPDB under code 26FG.

## References

1. Jain, R., Aggarwal, A. K. & Rechkoblit, O. Eukaryotic DNA polymerases. Curr. Opin. Struct. Biol. 53, 77–87 (2018).

2. Timinskas, K., Kazlauskas, D., Timinskas, A. & Venclovas, Č. Diversity and distribution of bacterial DNA polymerases. Nucleic Acids Res. 54, gkag133 (2026).

3. Raia, P., Delarue, M. & Sauguet, L. An updated structural classification of replicative DNA polymerases. Biochem. Soc. Trans. 47, 239–249 (2019).

4. Goodall, E. C. A. et al. The Essential Genome of *Escherichia coli* K-12. mBio 9, e02096–17 (2018).

5. Ghosh, S., Goldgur, Y. & Shuman, S. Mycobacterial DNA polymerase I: activities and crystal structures of the POL domain as apoenzyme and in complex with a DNA primer-template and of the full-length FEN/EXO–POL enzyme. Nucleic Acids Res. 48, 3165–3180 (2020).

6. Huberts, P. & Mizrahi, V. Cloning and sequence analysis of the gene encoding the DNA polymerase I from Mycobacterium tuberculosis. Gene 164, 133–136 (1995).

7. DeJesus, M. A. et al. Comprehensive Essentiality Analysis of the *Mycobacterium tuberculosis* Genome via Saturating Transposon Mutagenesis. mBio 8, e02133–16 (2017).

8. Gordhan, B. G., Andersen, S. J., De Meyer, A. R. & Mizrahi, V. Construction by homologous recombination and phenotypic characterization of a DNA polymerase domain polA mutant of Mycobacterium smegmatis. Gene 178, 125–130 (1996).

9. Brautigam, C. A., Sun, S., Piccirilli, J. A. & Steitz, T. A. Structures of Normal Single-Stranded DNA and Deoxyribo-3‘-*S* -phosphorothiolates Bound to the 3‘-5‘ Exonucleolytic Active Site of DNA Polymerase I from *Escherichia coli*,. Biochemistry 38, 696–704 (1999).

10. Rothwell, P. J. & Waksman, G. Structure and mechanism of DNA polymerases. in Advances in Protein Chemistry vol. 71 401–440 (Elsevier, 2005).

11. Toste Rêgo, A., Holding, A. N., Kent, H. & Lamers, M. H. Architecture of the Pol III– clamp–exonuclease complex reveals key roles of the exonuclease subunit in processive DNA synthesis and repair. EMBO J. 32, 1334–1343 (2013).

12. Sharma, A., Kottur, J., Narayanan, N. & Nair, D. T. A strategically located serine residue is critical for the mutator activity of DNA polymerase IV from Escherichia coli. Nucleic Acids Res. https://doi.org/10.1093/nar/gkt146 (2013) doi:10.1093/nar/gkt146.

13. Kottur, J. et al. Unique structural features in DNA polymerase IV enable efficient bypass of the N<sup>2</sup> adduct induced by the nitrofurazone antibiotic. Structure 23, (2015).

14. Johnson, M. K., Kottur, J. & Nair, D. T. A polar filter in DNA polymerases prevents ribonucleotide incorporation. Nucleic Acids Res. https://doi.org/10.1093/nar/gkz792(2019) doi:10.1093/nar/gkz792.

15. Bunting, K. A., Roe, S. M. & Pearl, L. H. Structural basis for recruitment of translesion DNA polymerase Pol IV/DinB to the β clamp. EMBO J. 22, 5883–5892 (2003).

16. Fernandez-Leiro, R., Conrad, J., Scheres, S. H. & Lamers, M. H. cryo-EM structures of the E. coli replicative DNA polymerase reveal its dynamic interactions with the DNA sliding clamp, exonuclease and τ. eLife 4, e11134 (2015).

17. Patoli, A. A., Winter, J. A. & Bunting, K. A. The UmuC subunit of the E. coli DNA polymerase V shows a unique interaction with the β-clamp processivity factor. BMC Struct. Biol. 13, 12 (2013).

18. Wijffels, G. et al. Inhibition of Protein Interactions with the β_2_ Sliding Clamp of *Escherichia coli* DNA Polymerase III by Peptides from β_2_ -Binding Proteins. Biochemistry 43, 5661– 5671 (2004).

19. Simonsen, S., Søgaard, C. K., Olsen, J. G., Otterlei, M. & Kragelund, B. B. The bacterial DNA sliding clamp, β-clamp: structure, interactions, dynamics and drug discovery. Cell. Mol. Life Sci. 81, 245 (2024).

20. Nirwal, S. et al. The structure of the MutL-CTD:processivity-clamp complex provides insight regarding strand discrimination in non-methyl-directed DNA mismatch repair. Nucleic Acids Res. 53, (2025).

21. López De Saro, F. J. & O’Donnell, M. Interaction of the β sliding clamp with MutS, ligase, and DNA polymerase I. Proc. Natl. Acad. Sci. 98, 8376–8380 (2001).

22. Bhardwaj, A., Ghose, D., Thakur, K. G. & Dutta, D. Escherichia coli β-clamp slows down DNA polymerase I dependent nick translation while accelerating ligation. PLOS ONE 13, e0199559 (2018).

23. Dalrymple, B. P., Kongsuwan, K., Wijffels, G., Dixon, N. E. & Jennings, P. A. A universal protein–protein interaction motif in the eubacterial DNA replication and repair systems. Proc. Natl. Acad. Sci. 98, 11627–11632 (2001).

24. Liebschner, D. et al. Polder maps: improving OMIT maps by excluding bulk solvent. Acta Crystallogr. Sect. Struct. Biol. 73, 148–157 (2017).

25. Letunic, I. & Bork, P. Interactive Tree of Life (iTOL) v6: recent updates to the phylogenetic tree display and annotation tool. Nucleic Acids Res. 52, W78–W82 (2024).

26. Lopez De Saro, F. J. Competitive processivity-clamp usage by DNA polymerases during DNA replication and repair. EMBO J. 22, 6408–6418 (2003).

27. Georgescu, R. E. et al. Structure of a small-molecule inhibitor of a DNA polymerase sliding clamp. Proc. Natl. Acad. Sci. 105, 11116–11121 (2008).

28. Bunting, K. A. Structural basis for recruitment of translesion DNA polymerase Pol IV/DinB to the -clamp. EMBO J. 22, 5883–5892 (2003).

29. Beuning, P. J., Simon, S. M., Godoy, V. G., Jarosz, D. F. & Walker, G. C. Characterization of Escherichia coli Translesion Synthesis Polymerases and Their Accessory Factors. in Methods in Enzymology vol. 408 318–340 (Elsevier, 2006).

30. Wolff, P. et al. Differential Modes of Peptide Binding onto Replicative Sliding Clamps from Various Bacterial Origins. J. Med. Chem. 57, 7565–7576 (2014).

31. Ditse, Z., Lamers, M. H. & Warner, D. F. DNA Replication in *Mycobacterium tuberculosis*. Microbiol. Spectr. 5, 5.2.20 (2017).

32. Kurthkoti, K. & Varshney, U. Distinct mechanisms of DNA repair in mycobacteria and their implications in attenuation of the pathogen growth. Mech. Ageing Dev. 133, 138–146 (2012).

33. Indiani, C., McInerney, P., Georgescu, R., Goodman, M. F. & O’Donnell, M. A sliding-clamp toolbelt binds high-and low-fidelity DNA polymerases simultaneously. Mol Cell 19, 805–815 (2005).

34. Castañeda-García, A. et al. A non-canonical mismatch repair pathway in prokaryotes. Nat. Commun. 8, 14246 (2017).

35. Ishino, S. et al. Activation of the mismatch-specific endonuclease EndoMS/NucS by the replication clamp is required for high fidelity DNA replication. Nucleic Acids Res. 46, 6206– 6217 (2018).

36. Takemoto, N., Numata, I., Su’etsugu, M. & Miyoshi-Akiyama, T. Bacterial EndoMS/NucS acts as a clamp-mediated mismatch endonuclease to prevent asymmetric accumulation of replication errors. Nucleic Acids Res. 46, 6152–6165 (2018).

37. Choe, K. N. & Moldovan, G.-L. Forging Ahead through Darkness: PCNA, Still the Principal Conductor at the Replication Fork. Mol. Cell 65, 380–392 (2017).

38. Acharya, N., Klassen, R., Johnson, R. E., Prakash, L. & Prakash, S. PCNA binding domains in all three subunits of yeast DNA polymerase δ modulate its function in DNA replication. Proc. Natl. Acad. Sci. U. S. A. 108, 17927–17932 (2011).

39. Wijffels, G. et al. Binding Inhibitors of the Bacterial Sliding Clamp by Design. J. Med. Chem. 54, 4831–4838 (2011).

40. Kling, A. et al. Targeting DnaN for tuberculosis therapy using novel griselimycins. Science https://doi.org/10.1126/science.aaa4690 (2015) doi:10.1126/science.aaa4690.

41. Pandey, P., Verma, V., Dhar, S. K. & Gourinath, S. Screening of E. Coli β-clamp inhibitors revealed that few inhibit Helicobacter pylori more effectively: Structural and functional characterization. Antibiotics https://doi.org/10.3390/antibiotics7010005 (2018) doi:10.3390/antibiotics7010005.

42. Paramythiotou, E. et al. A life-threatening case of disseminated nocardiosis due to *Nocardia brasiliensis*. Indian J. Crit. Care Med. 16, 234–237 (2012).

43. Li, T. et al. Gordonia bronchialis: an emerging opportunistic pathogen—a case report and comprehensive review. BMC Infect. Dis. 25, 1312 (2025).

44. Sharma, M., Narayanan, N. & Nair, D. T. The proofreading activity of Pfprex from Plasmodium falciparum can prevent mutagenesis of the apicoplast genome by oxidized nucleotides. Sci. Rep. 10, 11157 (2020).

45. Nirwal, S., Kulkarni, D. S., Sharma, A., Rao, D. N. & Nair, D. T. Mechanism of formation of a toroid around DNA by the mismatch sensor protein. Nucleic Acids Res. 46, 256–266 (2018).

46. Punjani, A., Rubinstein, J. L., Fleet, D. J. & Brubaker, M. A. cryoSPARC: algorithms for rapid unsupervised cryo-EM structure determination. Nat. Methods 14, 290–296 (2017).

47. Rubinstein, J. L. & Brubaker, M. A. Alignment of cryo-EM movies of individual particles by optimization of image translations. J. Struct. Biol. 192, 188–195 (2015).

48. Zivanov, J., Nakane, T. & Scheres, S. H. W. Estimation of high-order aberrations and anisotropic magnification from cryo-EM data sets in *RELION* -3.1. IUCrJ 7, 253–267 (2020).

49. Stagg, S. M., Noble, A. J., Spilman, M. & Chapman, M. S. ResLog plots as an empirical metric of the quality of cryo-EM reconstructions. J. Struct. Biol. 185, 418–426 (2014).

50. Punjani, A., Zhang, H. & Fleet, D. J. Non-uniform refinement: adaptive regularization improves single-particle cryo-EM reconstruction. Nat. Methods 17, 1214–1221 (2020).

51. Zivanov, J., Nakane, T. & Scheres, S. H. W. A Bayesian approach to beam-induced motion correction in cryo-EM single-particle analysis. IUCrJ 6, 5–17 (2019).

52. Waterhouse, A. et al. SWISS-MODEL: homology modelling of protein structures and complexes. Nucleic Acids Res. 46, W296–W303 (2018).

53. Punjani, A. & Fleet, D. J. 3DFlex: determining structure and motion of flexible proteins from cryo-EM. Nat. Methods 20, 860–870 (2023).

54. Pettersen, E. F. et al. UCSF Chimera—A visualization system for exploratory research and analysis. J. Comput. Chem. 25, 1605–1612 (2004).

55. Pintilie, G. D., Zhang, J., Goddard, T. D., Chiu, W. & Gossard, D. C. Quantitative analysis of cryo-EM density map segmentation by watershed and scale-space filtering, and fitting of structures by alignment to regions. J. Struct. Biol. 170, 427–438 (2010).

56. Zhang, H. et al. CryoPROS: Correcting misalignment caused by preferred orientation using AI-generated auxiliary particles. Nat. Commun. 16, 4565 (2025).

57. Kimanius, D., Dong, L., Sharov, G., Nakane, T. & Scheres, S. H. W. New tools for automated cryo-EM single-particle analysis in RELION-4.0. Biochem. J. 478, 4169–4185 (2021).

58. Zheng, S. Q. et al. MotionCor2: anisotropic correction of beam-induced motion for improved cryo-electron microscopy. Nat. Methods 14, 331–332 (2017).

59. Rohou, A. & Grigorieff, N. CTFFIND4: Fast and accurate defocus estimation from electron micrographs. J. Struct. Biol. 192, 216–221 (2015).

60. Scheres, S. H. W. Classification of Structural Heterogeneity by Maximum-Likelihood Methods. in Methods in Enzymology vol. 482 295–320 (Elsevier, 2010).

61. Cao, H. et al. EMReady2: improvement of cryo-EM and cryo-ET maps by local quality-aware deep learning with Mamba. Nat. Commun. https://doi.org/10.1038/s41467-026-71794-1 (2026) doi:10.1038/s41467-026-71794-1.

62. Tan, Y. Z. et al. Addressing preferred specimen orientation in single-particle cryo-EM through tilting. Nat. Methods 14, 793–796 (2017).

63. Meng, E. C. et al. UCSF CHIMERAX : Tools for structure building and analysis. Protein Sci. 32, e4792 (2023).

64. Emsley, P., Lohkamp, B., Scott, W. G. & Cowtan, K. Features and development of *Coot*. Acta Crystallogr. D Biol. Crystallogr. 66, 486–501 (2010).

65. Croll, T. I. *ISOLDE* : a physically realistic environment for model building into low-resolution electron-density maps. Acta Crystallogr. Sect. Struct. Biol. 74, 519–530 (2018).

66. Liebschner, D. et al. Macromolecular structure determination using X-rays, neutrons and electrons: recent developments in *Phenix*. Acta Crystallogr. Sect. Struct. Biol. 75, 861–877 (2019).

67. Kumar, A. et al. Protein crystallography beamline (PX-BL21) at Indus-2 synchrotron. J. Synchrotron Radiat. 23, 629–634 (2016).

68. Rösner, B. et al. Sharper, smaller, brighter: enhanced optical performance at the X06DA-PXIII beamline after the SLS 2.0 upgrade. J. Synchrotron Radiat. 33, 580–584 (2026).

69. Battye, T. G. G., Kontogiannis, L., Johnson, O., Powell, H. R. & Leslie, A. G. W. *iMOSFLM* : a new graphical interface for diffraction-image processing with *MOSFLM*. Acta Crystallogr. D Biol. Crystallogr. 67, 271–281 (2011).

70. Evans, P. R. & Murshudov, G. N. How good are my data and what is the resolution? Acta Crystallogr. D Biol. Crystallogr. 69, 1204–1214 (2013).

71. McCoy, A. J. et al. *Phaser* crystallographic software. J. Appl. Crystallogr. 40, 658–674 (2007).

72. Agirre, J. et al. The *CCP* 4 suite: integrative software for macromolecular crystallography. Acta Crystallogr. Sect. Struct. Biol. 79, 449–461 (2023).

73. Sayers, E. W. et al. Database resources of the National Center for Biotechnology Information in 2025. Nucleic Acids Res. 53, D20–D29 (2025).

74. Madeira, F. et al. The EMBL-EBI Job Dispatcher sequence analysis tools framework in 2024. Nucleic Acids Res. 52, W521–W525 (2024).

75. Crooks, G. E., Hon, G., Chandonia, J.-M. & Brenner, S. E. WebLogo: A Sequence Logo Generator: Figure 1. Genome Res. 14, 1188–1190 (2004).

76. Eisenberg, D., Schwarz, E., Komaromy, M. & Wall, R. Analysis of membrane and surface protein sequences with the hydrophobic moment plot. J. Mol. Biol. 179, 125–142 (1984).

