## Supplementary Figures and Legends for "Structural basis of interaction between Pol1 and DnaN from *Mycobacterium tuberculosis*"

**Supplementary figure legends**

**Figure S1: Assessment of the interaction of different constructs of Ec-PolI with Ec-β-clamp** (A) Micro-Scale Thermophoresis was used to measure the interaction of Ec-β-clamp with Ec-PolI-FL (red) and Ec-PolI-Klenow (blue). (B) Gel filtration analysis was carried out for Ec-β-clamp (red), Ec-PolI-FL (purple) and Ec-β-clamp mixed with Ec-PolI-FL (orange). The corresponding SDS-PAGE analysis of the eluted fractions from the sample containing a mixture of Ec-β-clamp and Ec-PolI-FL is shown. (C) Gel filtration analysis was carried out on Ec-β-clamp (red), Ec-PolI-Klenow (green) and Ec-β-clamp mixed with Ec-PolI-Klenow (dark blue). The corresponding SDS-PAGE analysis of the eluted fractions from the sample containing a mixture of Ec-β-clamp and Ec-PolI-Klenow is shown.

**Figure S2: 3DFlex workflow for structural heterogeneity analysis:** Schematic representation of the 3DFlex workflow for structural heterogeneity analysis and movie generation to find conformational flexibility in the complex.

**Figure S3: Cryo-EM data processing workflow for single particle analysis of Mtb-PolI-Klenow:Mtb-β-clamp complex.** Schematic diagram showing cryo-EM data processing workflow using Relion4.0. The particle subsets obtained after 3D classification from two independent datasets were processed separately using cryoPROS.

**Figure S4: Conservation analysis of CBM interacting residues of β-clamp from the members of the order Mycobacteriales:** MSA of β-clamp sequences from representative members of the Mycobacteriales order showed that the residues involved in interaction with the PolI CBM are conserved. The interacting residues are highlighted in cyan colour in the MSA.

**Figure S5: Conservation analysis of the CBM in PolI from pathogenic members of Mycobacteriales order:** Multiple sequence alignment of PolI sequences from pathogenic members of Mycobacteriales order shows that the QLSLLDD stretch (highlighted in green) is conserved.

**Supplementary Movie legends**

**Supplementary Movie S1: Dynamic interaction of the thumb domain of Mtb-PolI-Klenow with Mtb-β-clamp.** The movie generated from continuous heterogeneity analysis using 3DFlex of the 7.87 Å cryo-EM reconstruction reveals the dynamic nature of the interaction between the thumb domain of Mtb-PolI-Klenow and the non-canonical site on the second monomer of the Mtb-β-clamp dimer.

**Supplementary Movie S2: Electron potential map of 4.9 Å resolution of the Mtb-PolI-Klenow:Mtb-β-clamp complex.** The movie shows the improved electron potential map of the Mtb-PolI-Klenow:Mtb-β-clamp complex, generated from cryoPROS-assisted processing of cryoEM data.

**Supplementary Movie S3: Docking of component proteins into the 4.9 Å map.** The movie shows the crystal structure of Mtb-β-clamp (3RB9) and the homology model of Mtb-PolI-Klenow, docked into the electron density map, with a resolution of 4.9 Å.

**Supplementary Movie S4: Interaction of Mtb-β-clamp with CBP12.** The movie shows the surface representation of the Mtb-β-clamp peptide binding site, coloured according to the Eisenberg hydrophobicity scale, with the residues of the CBP12 peptide shown in stick representation and coloured according to atom.

### Figure S1

**A.**

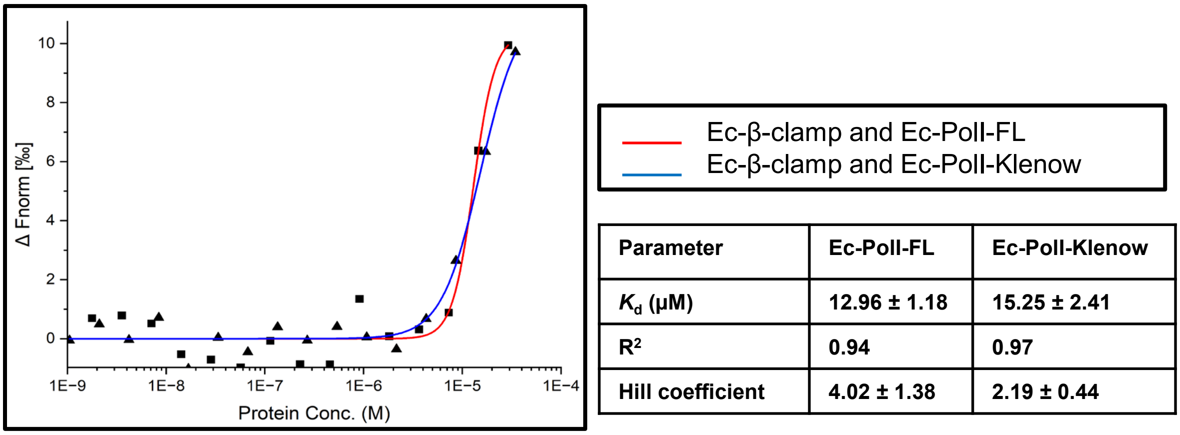


| Parameter | Ec-PolI-FL | Ec-PolI-Klenow |
|---|---|---|
| $K_d$ (μM) | 12.96 ± 1.18 | 15.25 ± 2.41 |
| $R^2$ | 0.94 | 0.97 |
| Hill coefficient | 4.02 ± 1.38 | 2.19 ± 0.44 |

**B.**

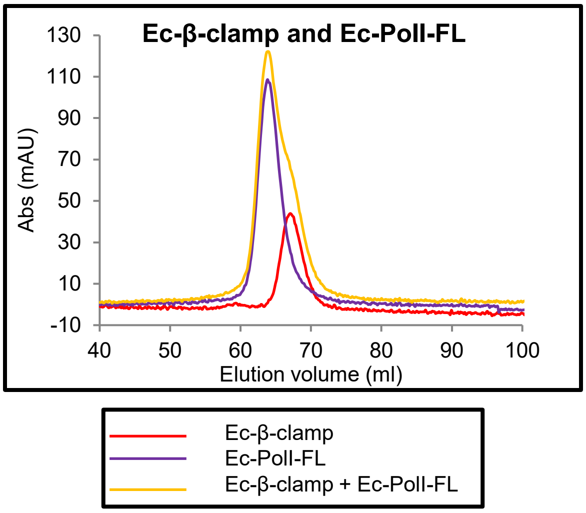


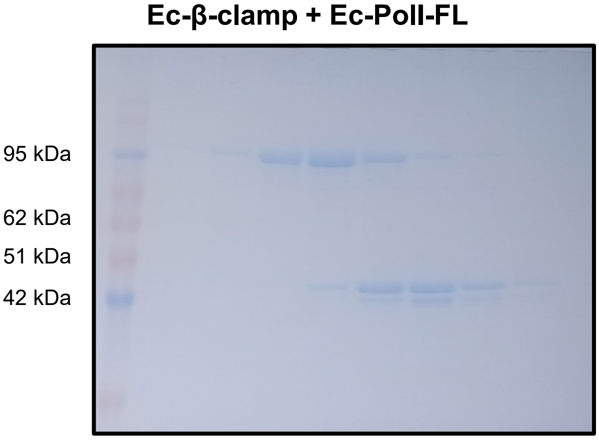


**C.**

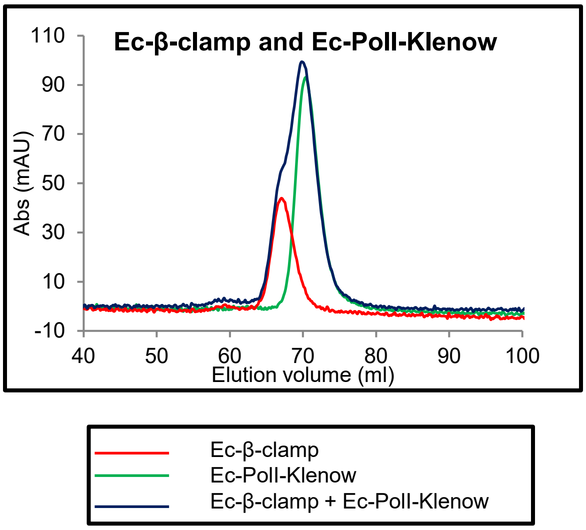


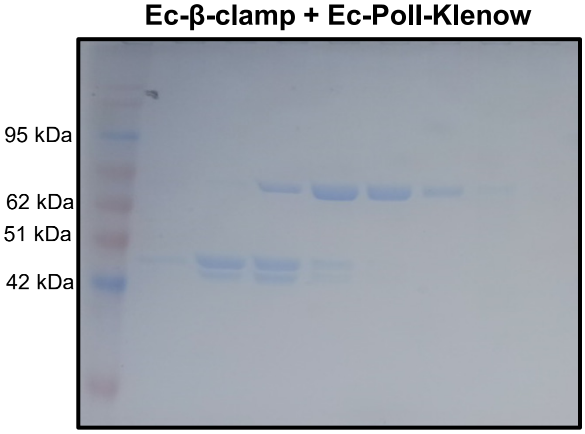

Figure S2

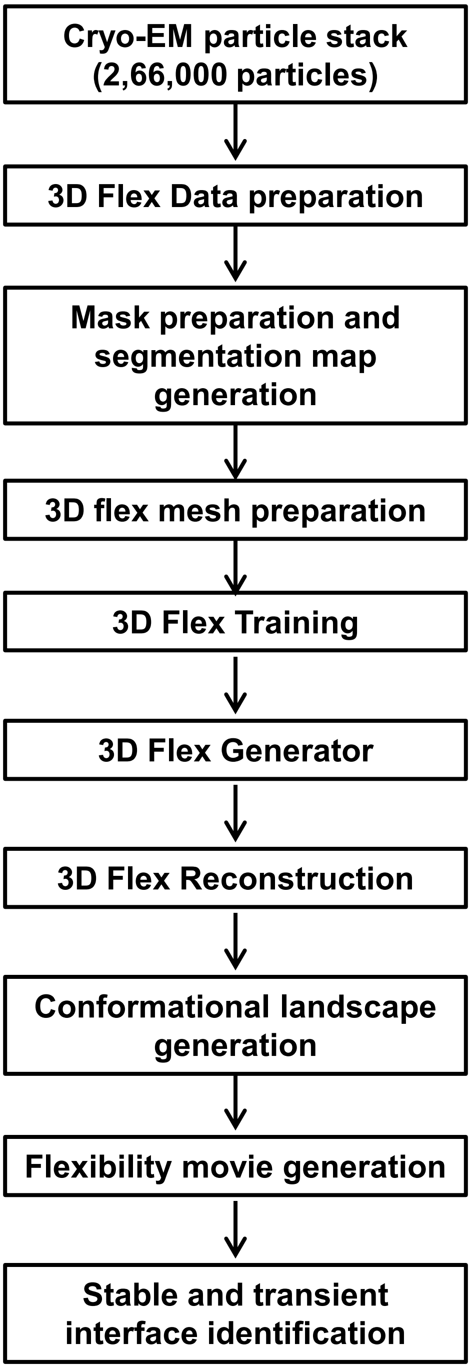

Cryo-EM particle stack (2,66,000 particles)
3D Flex Data preparation
Mask preparation and segmentation map generation
3D flex mesh preparation
3D Flex Training
3D Flex Generator
3D Flex Reconstruction
Conformational landscape generation
Flexibility movie generation
Stable and transient interface identification

### Figure S3

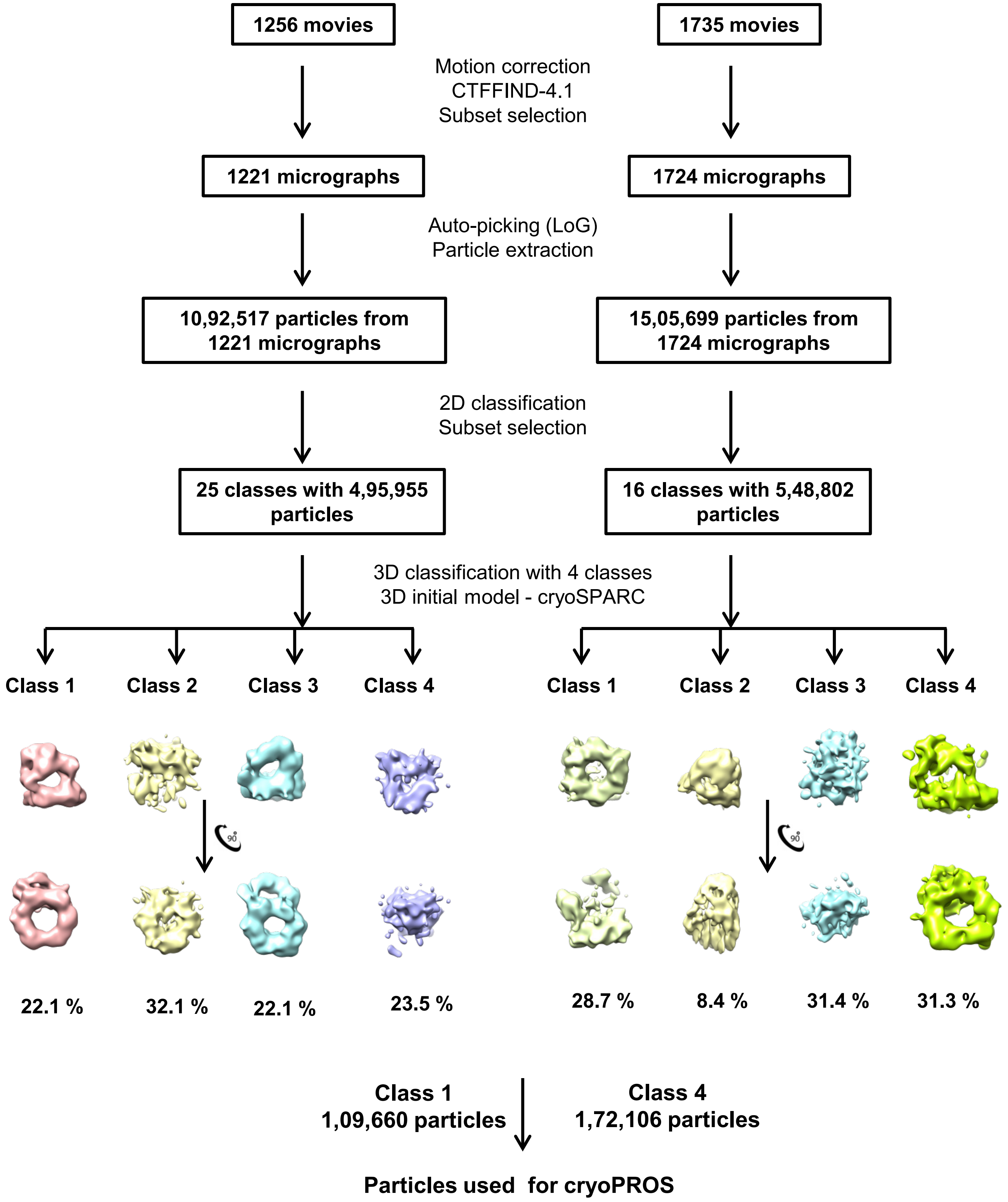
1256 movies
1735 movies
Motion correction
CTFFIND-4.1
Subset selection
1221 micrographs
1724 micrographs
Auto-picking (LoG)
Particle extraction
10,92,517 particles from 1221 micrographs
15,05,699 particles from 1724 micrographs
2D classification
Subset selection
25 classes with 4,95,955 particles
16 classes with 5,48,802 particles
3D classification with 4 classes
3D initial model - cryoSPARC
Class 1
Class 2
Class 3
Class 4
Class 1
Class 2
Class 3
Class 4
22.1 %
32.1 %
22.1 %
23.5 %
28.7 %
8.4 %
31.4 %
31.3 %
Class 1
1,09,660 particles
Class 4
1,72,106 particles
Particles used for cryoPROS

### Figure S4

```
M. tuberculosis     LAATDRFRLAVRELKWSASS-PDIEAAVLVPAKTLAEAAKAGI--GGSDVRLSLGT-GPG
N. halotolerans     LAATDRFRLAVRHLQWQPNR-SDIETAVLIPARTLSEAAKTLGSAS-APVQLSL---GNG
G. desulfuricans    LAATDRFRLAVRELEWEPEQ-ADINGAVLVPAKTLSESAKTAGAEGTGAVRLAFGS-GDG
T. sputi            LAATDRFRLAVREFEWQPAS-SDVKAAVLVPAKTLSEAAKTFAAD--PEVGIALGA-GSA
S. rugosus          LAATDRFRLSVRKLAWEGKDFADEPVTALVPAAVLSEFARTPLPSDDTPVEFALDL---G
C. ammoniagenes     LTATDRFRLAVRTFEWEPTA-EDVKTQLLVPAKTLQETARTLDSHINDPIEIAVGT-GDS
S. jeojiensis       LAATDRFRLAVRELEWMPTN-GETNTAVLVPARTLSESAKTLGGGANSPVELALGS-GSN
H. rhizosphaerae    LAATDRFRLAVRSIDWIPEK-ADTEISVLVPAKTLSEAGKSVGGSGNSIVKIALGE-GHS
D. aurantiaca       LIATDRFRLAIREFEWEPAR-DDVAVEVLIPAKALSEVTRSAGAG--GRVDLSLGA-GSE
L. clevelandensis   FAATDRFRLAVRKFEWDPFA-SFSDAAVLIPNKMLSETAKTFAATSTAPVELALGGANDT
T. fengzijianii     MAATDRFRLAVREFEWKPAT-PELEAAVLVPARTLAEAAKTMTASASADIEIEMGD-GSS
                    : *******::* : *            *:*   * *  ::        : : .

M. tuberculosis     VGKDGLLGISGNGKRSTTRLLDAEFPKFRQLLPTEHTAVATMDVAELIEAIKLVALVADR
N. halotolerans     GGGDGLLGIVNAGRRTTTRLLDAEFPKFRQLLPKEHTSIATLPVSVLSEAIKRVALVAER
G. desulfuricans    IGAEGILGIILGETKQTTTRLLDADFPKFRQLLPASHTAVATIESAPLIEAIKRVSLVAER
T. sputi            VGADGLLGVVGADRRTTTRLLDADFPKFRQLLPASHTAIATMEIAPLLEAIKRVALVADR
S. rugosus          SGSPSLLGVQAGPFRITTRLLDAQLPDVSQLIPKRYTAVARVDLAPLVEAVRRVSLVSAR
C. ammoniagenes     IAADGLFGLHADNRETTTRMLDADFPTVSHLLPKTHTAMASVEIGPLQEAIRRVSLLTDR
S. jeojiensis       IGADGLLGIVGQGRRTTTRLLDAEFPKFRQLLPNEHTALATVEVTPLVEAIKRVALVAER
H. rhizosphaerae    LASEGLLGIISSTRRTTSRLLDAEFPKFRQLLPKTHTAMAALEVASLIEAIKRVALLAER
D. aurantiaca       VGAEGILGVLVSGQRTTTRLLDAEFPKVRQLLPPQHNSIAVVEVDALIQAIKRVALVADR
L. clevelandensis   LGGEGLLGIVGENRRITTRLLDAQFPPFRTLLPKAHNAIATVDIASLVDSIKRVSLMSDR
T. fengzijianii     VGAEGMLGVTSGPRQTTARLLDAEFPKFRQLLPATHTAVAAVEVAPLTEAIKRVALVAER
                     .  .::*:     . *:*:***::* .  *:*  :.::* :    * :::: *:*:: *

M. tuberculosis     GAQVRMEFADG----SVRLSAGADDVGRAEEDLVVDYAGE-PLTIAFNPTYLTDGLSSLR
N. halotolerans     GAQVRLEFSTD----GVLLSAGGDDAGRAEEWLEADFRGE-PLTIAFNPGYLTEGLAALH
G. desulfuricans    GAQVRMEFSGG----SVLLTAGGDEAGKAEEELPVEFQGE-PLTIAFNPGYLQDGLSAIN
T. sputi            GAQVRMEFADG----TLRLSAGGDDAGKAEEELPADFQGE-PLTIAFNPGYLQDGLAAVH
S. rugosus          RSQIRLDFESSGEDSVLRLSAGGEEAGFAEEELPVRHWGE-SISVVFNPGYLLDGLSTFR
C. ammoniagenes     NAQIRMLFSEG----EVQLAAGATDTGNAEETIPCAFTGRDELLIAFNPAFLKDGLAVIP
S. jeojiensis       GAQVRMEFSAE----GLLLSAGGDDAGRAEESLPAEFQGE-PLIIAFNPGYLLDGLGAMR
H. rhizosphaerae    GAQIRMDFSHD----GLQISAGGDDAGQAEETLPIEFRGE-PLVIAFNPGYLIDGLNSLH
D. aurantiaca       GVQVRLAFNEG----ELALSAGGDDAAQANETLPVDFVGE-PLTIAFNPGYLLDGLGSVH
L. clevelandensis   GAQVRLAFTPE----SVTLTAGGDDSGEAEETLPVKYWGE-PMTIAFNPSYLLDGLSVLD
T. fengzijianii     GAQVRMQFSED----GLVLAAGGDDSSSAEESLPAEFAGE-PLTIAFNPGYLQEGLGAVG
                      *:*: *        : ::**. : . *:* :   . *.  : :.*** :* :**  .

M. tuberculosis     SERVSFGFTTAGKPALLRPVSGDDRPVAGLNGNGPFPAVSTDYVYLLMPVRLPG
N. halotolerans     TDRVTFGFTTPSRPAVLLPASDEEPKV---LESGAFAALESQYIYLLMPVRLPG
G. desulfuricans    AKSVDFGFTTPSRPAVLRPSAGERPVP---DESGAFVAPESVFSYLLMPVRLPG
T. sputi            TESVTFGFTTPSRPAVLRPAVDNGYEP---DDSGAFPAPQSPYTYLLMPVRLPG
S. rugosus          AEQAQFGFTGQNRAVALVSGDGAEPEA----IAEGYTAGSSDQVYVLMPVRLQG
C. ammoniagenes     TDRVVFGFTEPSRPAIMIPEPEELPEA---DEDGNFPTPDTFFTYLLMPVRLPG
S. jeojiensis       SERALFGFTNPSRPAVLRPAGDEPLSP---DESGAFAAPSSDYTYLLMPVRLPG
H. rhizosphaerae    TDKVVFGFTLPSRPAVLQPADSDVELP--DGEDGAFAARDSEYTYLLMPVRLPG
D. aurantiaca       SSRVAFGFTQASRPAVLRPAPETLPEP---DADGAIAPVDSNHTYLLMPVRLPG
L. clevelandensis   GTTASFGFTTPNRPAVIIPGIDNEPEA---NDAGEYPAPDTDYIYLLMPVRLPG
T. fengzijianii     TEHVHLGFTSPSRPAVLCPAAEEFPQA---DDSGAFPASSGSYTYLLMPVRLPG
                       . :***  .: . :                      .    *:****** *
```

### Figure S5

```
M. tuberculosis      -QQQQLSLLDDDD-TDAETIQTTILRARAVIDLADA
M. bovis             -QQQQLSLLDDDD-TDAETIQTTILRARAVIDLADA
M. africanum         -QQQQLSLLDDDD-TDAETIQTTILRARAVIDLADA
M. microti           -QQQQLSLLDDDD-TDAETIQTTILRARAVIDLADA
M. canettii          -QQQQLSLLDDDD-TDAETIQTTILRARAVIDLADA
M. pinnipedii        -QQQQLSLLDDDD-TDAETIQTTILRARAVIDLADA
M. caprae            -QQQQLSLLDDDD-TDAETIQTTILRARAVIDLADA
M. leprae            -EQEQFSLLDNVDEVDKQAIQTLILRARAVVDLAAA
M. lepromatosis      -EQEQFSLLDNVDEVDKQTVQTLILRARAVADLAAA
M. avium             -QQQQLSLLDDTDGTDDQAVQTLILRAVAVLDLADA
M. intracellulare    -EQQQLSLLDDTDGTDDQAVQTLLLRAVAVLDLADA
M. kansasii          -EQQQLSLLDDMEGVDDQAVQTAILRARAVADLAEA
M. marinum           -EQQQLSLLDDDD-TDVAAAQTAILRARAVIDLADA
M. ulcerans          -EQQQLSLLDDDD-TDVAAAQTAILRARAVIDLADA
M. scrofulaceum      -EQQQLSLLDDPEGTDDQAVQTLILRAVAVLDLADA
M. xenopi            -EQQQLSLLDDTDGVDDQAVQTTILRARAVADLADA
M. malmoense         -EQQQLSLLDSTDGVDEQAVQTLILRAQAVVDLADA
M. abscessus         -GEQQLSLLDEDGAGDAKAAQAQMLRARAVIDLADA
M. fortuitum         -EQQQLSLLDDSDGVDEQAVQTVILRACAVLDLADA
M. chelonae          -GEQQLSLLDDDGAGDAKAAQAQMLRARAVIDLADA
N. asteroids         -DSGQLSLLDDEDTVAEQTARELMLRARAVADLADA
N. brasiliensis      -ETPQLSLLDDEDTVDAELAQTEMLRARAVADLADA
R. equi              -GQQQLSLLDDEDAADAEAAEGEILRSRAVRDLAAA
Gordonia             GGDGQLSLLDDDAD-TQRAADQLMLSAQAVAELADT
Tsukamurella         -DDSQLSLLDDPDDSAGEKAQQAILRARAVADLADA
                         *:****.            :* : ** :** :
```
